# How visual scenes recruit spatial maps: threshold-gated activation in CA3 versus continuous integration in CA1

**DOI:** 10.64898/2026.07.22.740220

**Authors:** Eun-Ho Lee, Inah Lee

## Abstract

The hippocampal cognitive map is constructed from environmental sensory inputs, but the conditions that recruit a spatial map from a quiescent baseline remain unclear. Using a parametric virtual reality paradigm with simultaneous recordings from CA1 and CA3 in rats, we demonstrate that these subregions follow fundamentally different rules. CA1 place cells exhibited robust selectivity under sparse conditions, continuously updating their tuning as landmarks accumulated. Conversely, CA3 remained quiescent, its recruitment nonlinearly gated until landmarks formed a spatiotemporally integrated ensemble, governed by ensemble density and configuration rather than absolute count. Naturalistic backgrounds facilitated early activation, eliciting stepwise recruitment as scene complexity increased. These transitions paralleled LFP dynamics shifting from sensory decoupling (reduced fast-gamma coupling) to global network engagement (increased slow-gamma coupling). Together, these findings establish a circuit-level framework wherein CA3 operates as a nonlinearly gated scene constructor and CA1 as a continuous landmark integrator to build and maintain the cognitive map.

## INTRODUCTION

The hippocampal ‘place cell’ is a key neural substrate of the cognitive map^1^, supported by subregional architectures with distinct computational roles. CA3 features recurrent collaterals forming an autoassociative network capable of nonlinear attractor dynamics^2,3^, whereas CA1 receives converging entorhinal cortex (EC) and CA3 inputs, hypothesized to act as a comparator^4–6^. These distinct architectures suggest that the emergence of spatial codes is governed by fundamentally different rules across subregions^7,8^—potentially segregated by fast and slow gamma oscillations routing feedforward EC inputs and internal CA3 processing, respectively^9^. Specifically, a new CA3 attractor state may require the coordinated activation of a neuronal ensemble to reach a critical threshold^10,11^, suggesting that the conditions triggering CA3 map formation differ qualitatively from those sufficient for CA1.

The conditions triggering hippocampal map formation can be conceptualized at three distinct levels: remapping, the reorganization of an already-established map in response to environmental changes^12,13^; formation, the biophysical mechanisms—such as behavioral timescale synaptic plasticity (BTSP)—enabling individual neurons to acquire spatial tuning^14–17^ and recruitment, the macroscopic transition of a functionally quiescent neural population—encompassing both silent and non-spatial active neurons—into a functional spatial representation^18,19^. Of these, recruitment is the least understood. While remapping and place-field formation are well characterized, the environmental conditions that initiate network-level map formation from a quiescent baseline—what we term *’recruitment conditions*’—remain poorly defined. Specifically, it is unknown whether this transition reflects passive sensory aggregation or a nonlinearly gated network state transition. This distinction matters: if CA3 requires coordinated ensemble inputs to cross an activation threshold, its recruitment would be fundamentally discontinuous, whereas CA1 may incrementally integrate any level of sensory input.

Visual stimuli prominently shape hippocampal spatial coding^12,20–22^, but prior studies utilizing highly variable visual inputs obscure which specific features are necessary and sufficient for map recruitment. A critical distinction lies between individual ‘landmarks’ and holistic ‘scenes.’ Landmarks function as independent, localized stimuli that support position identification^20,23^, whereas scenes represent more complex, spatially coherent ensembles of visual elements—encompassing object configurations and background features—that together define a global environmental context^24–26^. This distinction aligns with Scene Construction Theory (SCT), which proposes that a core hippocampal function is to construct and maintain such coherent internal models of the environment^27^, a capacity that extends to rodent scene-based navigation and memory^28–31^. However, because SCT has been supported primarily by human neuroimaging^24,32,33^, its circuit-level implementation remains unresolved. We hypothesize that this scene-landmark distinction corresponds directly to the differential recruitment rules of CA3 and CA1: while CA1 incrementally integrates individual landmarks, CA3 may require scene-like stimulus configurations—specifically, a spatially coherent ensemble—to cross the threshold for attractor state formation.

Our previous work provided initial evidence that CA3 is selectively recruited under visual ensemble conditions, whereas CA1 responds to sparse landmarks^34^. However, because those sensory manipulations were discontinuous, the precise quantitative thresholds and configural rules governing CA3 recruitment remained unresolved. Here, we directly test whether CA3 recruitment requires scene-like inputs by implementing a parametric virtual reality (VR) paradigm that continuously varies visual ensemble density, spatial configuration, and background scene complexity during simultaneous CA1 and CA3 recordings. This design reveals where recruitment shifts along these dimensions, and whether such transitions are accompanied by distinct network-level dynamics. Our results demonstrate that CA1 and CA3 follow fundamentally different rules for map recruitment, revealing the circuit-level dynamics by which the hippocampus balances continuous sensory integration with discrete scene construction.

## RESULTS

### CA1 and CA3 diverge in recruitment along a visual gradient

To determine whether hippocampal place cells encode the quantitative accumulation of visual landmarks through linear integration or nonlinear state transitions, we developed a parametric landmark-manipulation paradigm in VR. Body-restrained rats navigated a virtual linear track (Figure 1A) where visual landmarks were presented within a controlled environment partitioned into landmark-visible (LV) and landmark-invisible (LIV) zones (Figure 1B). By constraining the animals’ field of view (FOV), we ensured that spatial firing was driven predominantly by experimenter-defined visual cues. To implement a continuous sensory gradient, the number of landmarks within the LV zone was incrementally scaled from 2 to 20 within each session in both ascending and descending sequences (Figure 1C) and grouped into five discrete blocks. Rats exhibited stable and consistent running velocities across all landmark conditions and sequence directions, ensuring uniform behavioral engagement (Figure S1).

**Figure 1.**
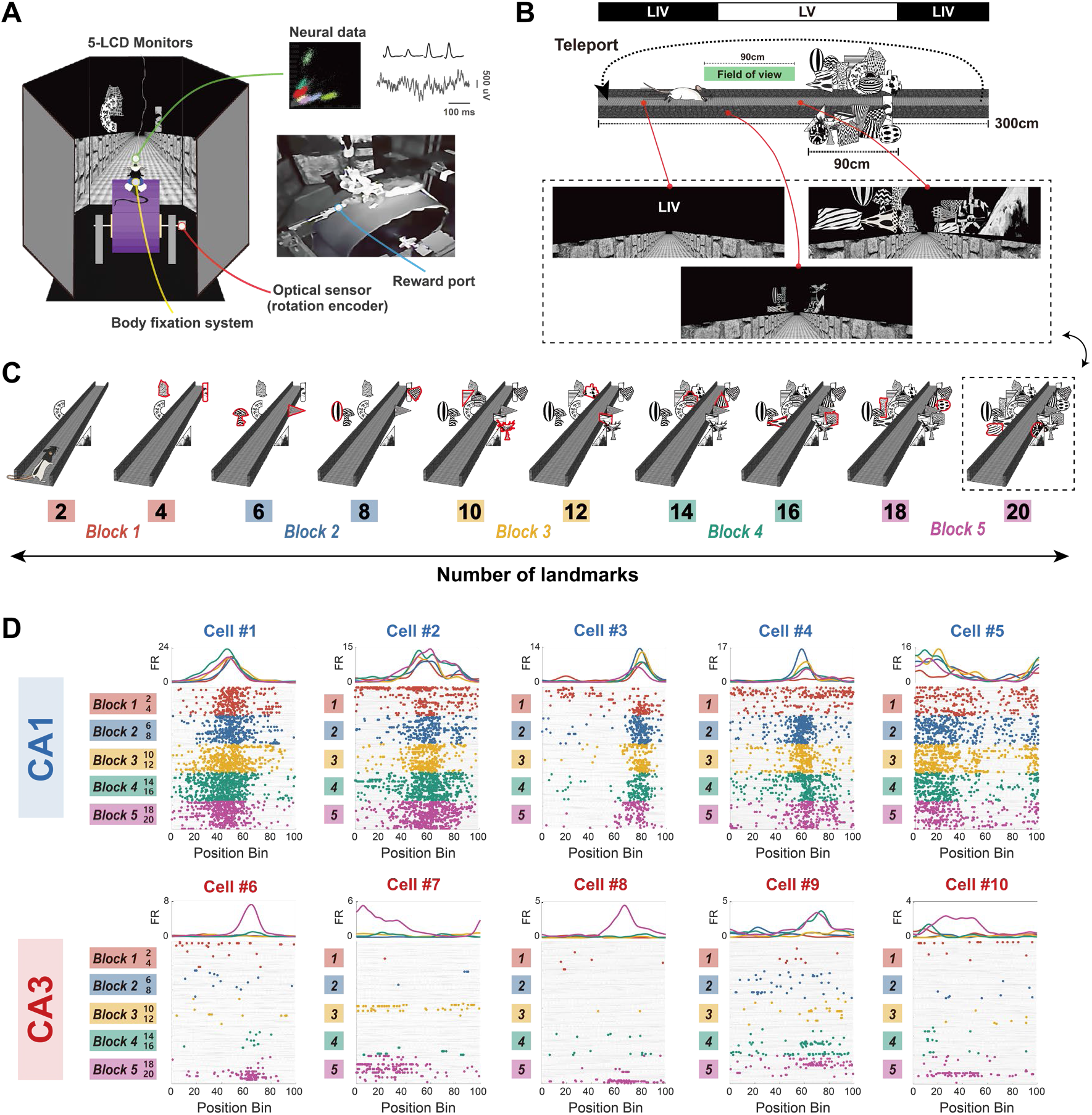
Parametric manipulation of the number of visual landmarks and differential response patterns of CA1 and CA3 place cells. (A) Schematic of the virtual reality (VR) experimental apparatus for body-restrained rats running on a cylindrical treadmill (n=8). (B) Configuration of the virtual linear track. The 300-cm track was divided into the landmark- visible (LV) zone (180 cm) and the landmark-invisible (LIV) zone (120 cm). Visual landmarks were distributed within a 90 cm segment of the LV zone. The rat’s field of view (FOV) was restricted to a 90-cm window of the track, ensuring that landmarks became visible only upon approach within the LV zone. Bottom panels illustrate the rat’s perspective at different locations on the track. (C) Experimental paradigm for the parametric manipulation of visual landmarks. The session comprised 10 sequential conditions. Initially, 2 landmarks were presented within the LV zone. Every 15 laps, two landmarks were added, incrementally increasing the total number in steps of two up to 20 (i.e., 2, 4, 6, …, 20). Red outlines indicate newly added landmarks for each condition. Note that all previously presented landmarks remained in place throughout the experiment. (D) Representative responses of CA1 (top, Cell #1-5) and CA3 (bottom, Cell #6-10) place cells across 5 blocks. Simultaneous recordings of single-unit activity were obtained from dorsal CA1 (n = 904 units) and CA3 (n = 815 units). Raster plots (left) showing spikes (colored dots) along the track for each block. Positional tuning curves (right) show the mean firing rate (Hz) along the track for each block. Note that CA1 cells form distinct place fields from early blocks, whereas CA3 cells exhibit abrupt place field formation, typically in later blocks.

Simultaneous CA1 and CA3 recordings revealed distinct activity patterns along this visual gradient. CA1 place cells exhibited robust spatial selectivity even under sparse landmark conditions (Block 1; 2-4 landmarks), maintaining stable place fields as additional landmarks were incorporated into an established spatial framework (Figure 1D). In contrast, CA3 units showed a nonlinear recruitment profile: most remained functionally quiescent during earlier blocks, with robust place fields emerging abruptly only at the highest landmark density (Block 5; 18–20 landmarks). Histological and longitudinal waveform analyses confirmed that this quiescence was not due to recording instability; CA3 units silent during navigation nonetheless fired robustly during sleep (Figure S2). Thus, CA3 quiescence under sparse landmark conditions reflects a genuine physiological response to landmark scarcity, demonstrating a fundamental divergence in how CA1 and CA3 transform sensory cues into spatial representations.

### CA1 generalizes while CA3 orthogonalizes across landmark blocks

To quantify these divergent response profiles at the population level, we assessed neuronal activity persistence across blocks. By tracking the number of blocks in which each active unit fired, we found that CA1 populations exhibited high persistence: a majority of units maintained activity across all five blocks (56.4%; χ²_(4)_ = 487.39, p < 0.0001; goodness-of-fit test), indicating CA1 recruits a stable neuronal pool to incrementally update a continuous spatial map (Figure 2A). In contrast, CA3 activity was highly transient; nearly half of active units (47.2%; χ²_(4)_ = 315.40, p < 0.0001) were recruited in only a single block. Overall persistence in CA3 was significantly lower than in CA1 (χ²_(4)_ = 378.75, p < 0.0001; χ² test of homogeneity).

**Figure 2.**
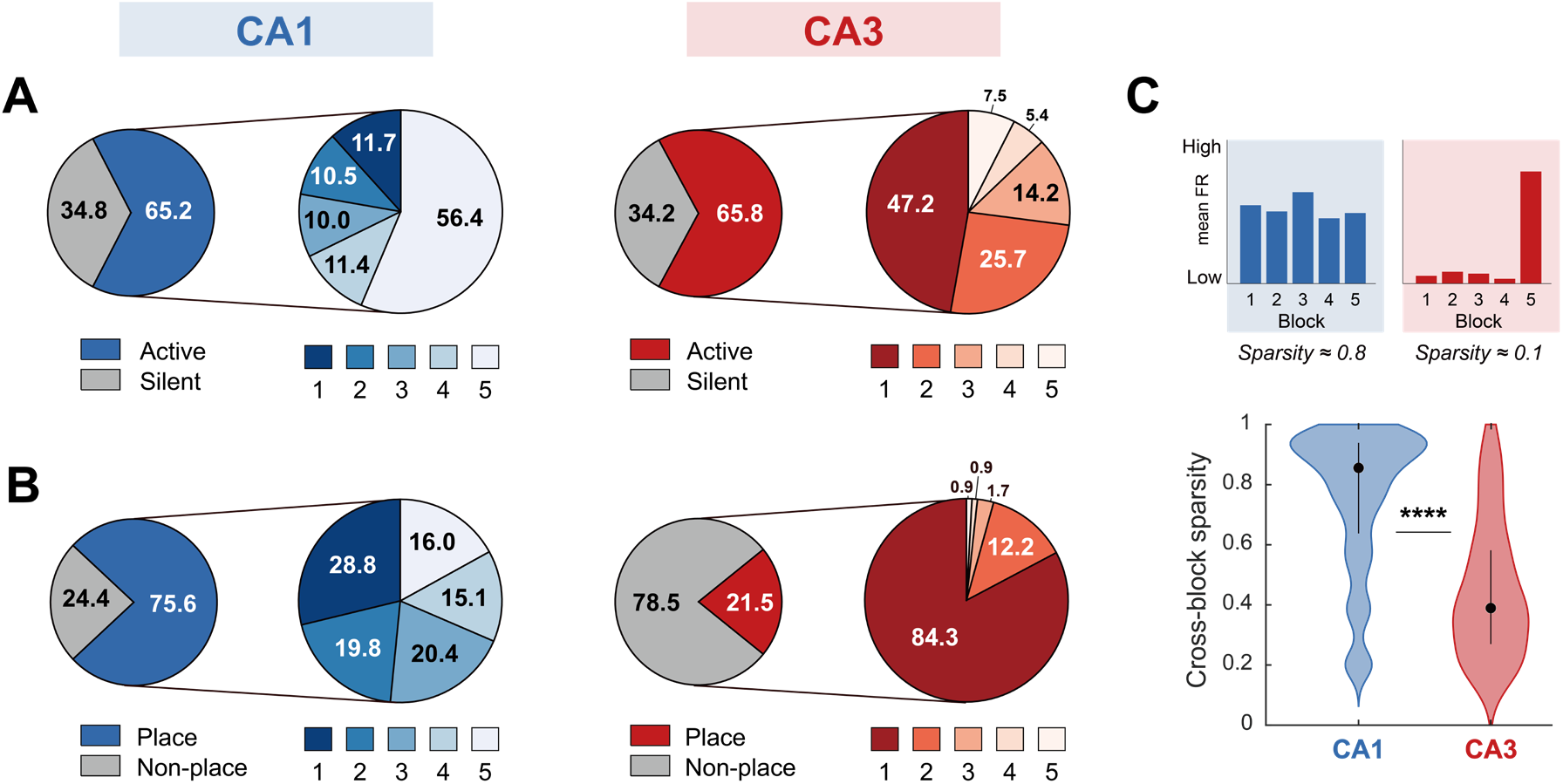
Distinct coding strategies across landmark conditions in CA1 and CA3. (A) Proportions of active cells (CA1, n = 589; CA3, n = 536) and their recruitment persistency. (left) Overall proportion of active cells. A cell was defined as ‘active cell’ if its mean firing rate exceeded 0.3 Hz in at least one of the 5 blocks. (right) Distribution of activity persistence. Segments represent the proportion of units active in N blocks. (B) Proportion of place cells and their block specificity. (left) Proportion of place cells within the active population. (right) Distribution of place field persistence across blocks. The segments represent the proportion of units identified as place cells in N blocks. (C) Comparison of cross-block sparsity between CA1 and CA3. Top: Conceptual illustration of the sparsity metric to distinguish persistent firing from orthogonalized activity; a value close to 1 indicates consistent firing across blocks (Dense), while a value close to 0 indicates activity restricted to a few blocks (Sparse). Bottom: Violin plots showing that CA3 cells have significantly lower cross-block sparsity values compared to CA1, indicating that CA3 representations are fundamentally more block-specific. **** p < 0.0001

This regional dissociation was more pronounced among place cells (Figure 2B). CA1 place cells frequently preserved their spatial fields across multiple blocks (71.2% recurring in ≥2 blocks; χ²_(4)_ = 26.23, p < 0.0001; goodness-of-fit test). Conversely, most CA3 place cells fired in a block-specific manner (84.3%; χ²_(4)_ = 302.87, p < 0.0001)—typically forming fields exclusively in Block 5—while remaining quiescent otherwise. The proportion of block-specific firing was significantly higher in CA3 than in CA1 (χ²(4) = 123.75, p < 0.0001; χ² test of homogeneity). To evaluate representational specificity, we computed a cross-block sparsity index, for which lower values indicate firing restricted to fewer blocks (Figure 2C). CA3 exhibited significantly lower sparsity than CA1 (Z = 20.69, p < 0.0001; Wilcoxon rank-sum test), confirming that CA3 recruits largely non-overlapping subpopulations to represent distinct visual configurations. Thus, CA1 generalizes a single neuronal pool across visual conditions whereas CA3 orthogonalizes them into non-overlapping subpopulations, reflecting two complementary coding strategies.

### CA3 undergoes a discrete, nonlinear state transition

We next examined the temporal dynamics of place-field formation as landmarks accumulated. Population rate maps revealed that CA1 generated a continuous spatial map from the outset (Block 1), whereas CA3 activity remained sparse and fragmented before undergoing an abrupt increase in place-cell recruitment—despite linear increases in environmental complexity (Figures 3A and S3).

**Figure 3.**
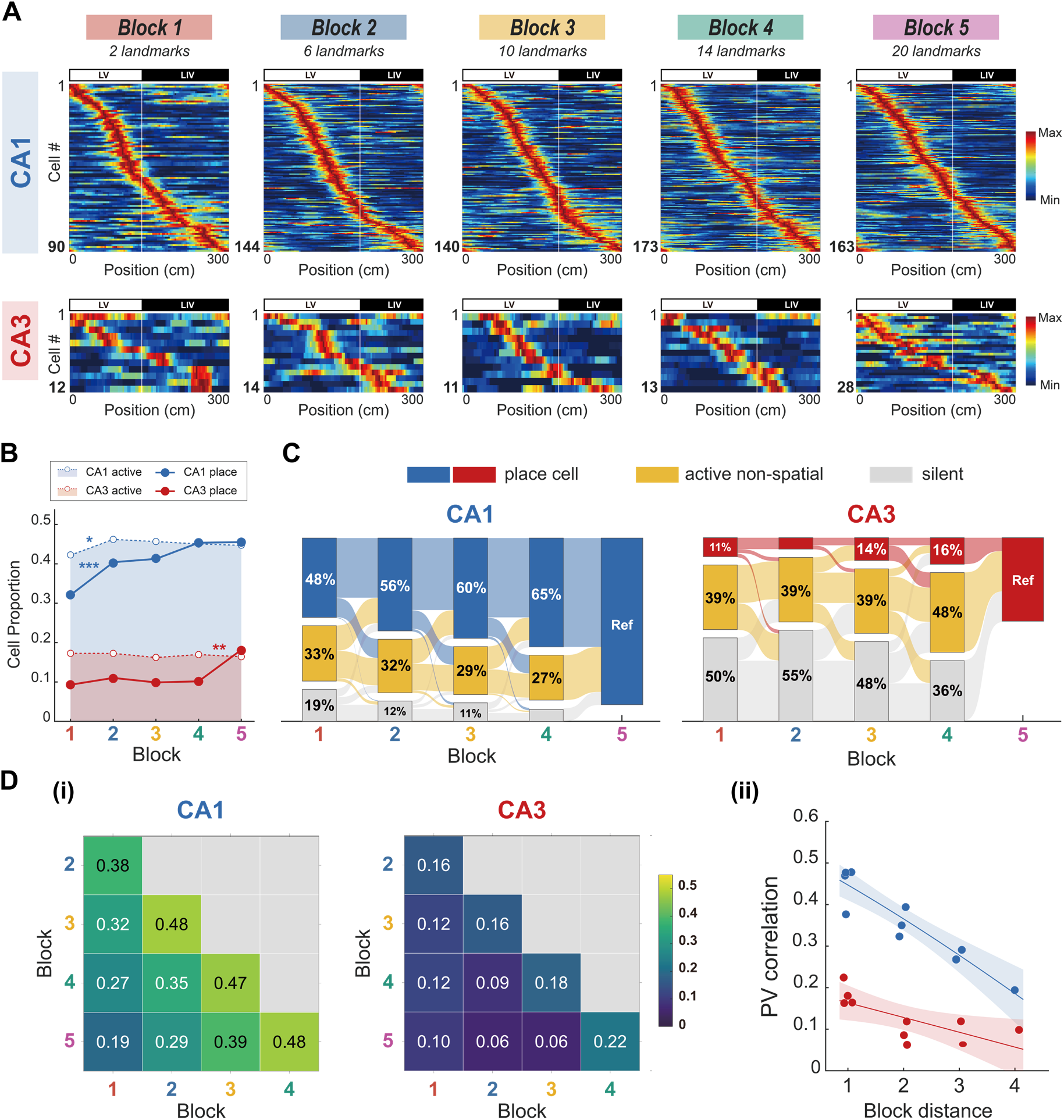
Divergent recruitment dynamics of CA1 and CA3 place cells in response to increased visual landmarks. (A) Population firing rate heatmaps for CA1 (top) and CA3 (bottom) across the 5 blocks. Each heatmap illustrates place cell activity from a single, representative landmark condition within its corresponding block. For visualization, the track position is aligned so that the LV zone (0- 180 cm) begins at 0 cm, followed by the LIV zone (180-300 cm). Each row represents a single place cell, sorted by its peak firing location; note the differences in the number of cells used to construct the population maps, indicated on each y-axis. (B) Proportion of active cells (open circles with dotted line) and place cells (filled circles with solid line) across blocks. The overall proportion of active cells remained stable across blocks in both regions (CA1: χ²(4) = 3.55, p = 0.47; CA3: χ²(4) = 2.35, p = 0.67; χ² tests). (C) Alluvial diagrams illustrating the retrospective state classification of units identified as place cells in Block 5 (Ref). Functional states in preceding blocks (1-4) are categorized as place cell (blue for CA1, red for CA3), active non-spatial (yellow), or silent (grey). (D) Population vector (PV) correlations from all place cells. (i) PV correlation matrices for CA1 (left) and CA3 (right); lower triangle only; with the diagonal masked and excluded from the color scale (0-0.55). (ii) PV correlation of every block pair (n = 10 per region) against block distance, with linear regression on Fisher-Z transformed coefficients and 95% confidence intervals. In CA3, a shallow negative slope reached significance (slope = −0.036, t(8) = −2.72, p = 0.026) but was confined to immediately adjacent blocks, consistent with the partial overlap of visual input between neighboring conditions rather than a graded drift of the map. * p < 0.05, ** p < 0.01, *** p < 0.001.

To distinguish changes in global excitability from changes in spatial selectivity, we compared the proportions of active and place cells across blocks. While overall active cell fractions remained stable in both regions, place-cell recruitment diverged significantly (Figure 3B). CA1 place-cell fractions increased gradually with landmark accumulation before saturating (among conditions: χ²_(4)_ = 37.18, p < 0.0001; Block 1 vs. 2: χ²_(1)_ = 11.22, p = 0.0008; χ² tests; Bonferroni-corrected α = 0.0125). In contrast, CA3 recruitment remained minimal through Blocks 1–4 and increased in a stepwise manner at Block 5 (among conditions: χ²_(4)_ = 17.23, p = 0.002; Block 4 vs. 5: χ²_(1)_ = 7.26, p = 0.007; α = 0.0125). Sigmoidal modeling confirmed this divergence: CA1 recruitment followed a relatively gradual curve with an early inflection (∼6 landmarks), whereas CA3 map emergence was characterized by a steep, late-stage transition (∼17 landmarks), confirming that its abruptness was robust across anatomical segments and criteria (Figure S4). This confirms that CA3 quiescence reflects a specific absence of spatial coding rather than global network suppression.

To identify the prior functional states giving rise to this recruitment, we tracked individual units retrospectively (Figure 3C). CA1 place cells identified in later blocks typically originated from already-active or spatially tuned units. In contrast, newly recruited CA3 place cells in Block 5 arose predominantly from previously quiescent units. This discrepancy in state distributions between the two subregions was highly significant from the earliest stage and persisted until the final transition phase (Block 1: χ²_(2)_ = 28.25, p < 0.0001; Block 4: χ²_(2)_ = 46.18, p < 0.0001; χ² tests). These findings demonstrate that the CA3 spatial map emerges through a discrete state transition—occurring independently of changes in single-unit firing rate or bursting properties (Figure S5A)—rather than the progressive refinement observed in CA1.

Population vector (PV) correlations confirmed this dissociation (Figure 3D). CA1 correlations decreased systematically with block distance (slope = -0.098, t_(8)_ = -7.1, p < 0.0001; linear regression on Fisher-Z transformed coefficients), reflecting a continuously updating map. Conversely, CA3 correlations remained uniformly low across all transitions, with only residual similarity between immediately adjacent blocks, indicating a highly orthogonalized representational structure. Overall correlations were significantly lower in CA3 than in CA1 (difference = -0.38, t_(16)_ = -8.85, p < 0.0001; GLM main effect of Region), and the decay slope in CA1 was significantly steeper than in CA3 (t_(16)_ = 3.23, p = 0.005; GLM Region × Distance interaction). Therefore, the abrupt recruitment in CA3 corresponds to the emergence of a distinct network state—marked by a concurrent increase in intra-regional synchronization (Figure S5B)—that is functionally different from the fragmented representations present during earlier blocks.

### Visual ensemble density, not landmark count, governs CA3 recruitment

CA3 recruitment in the accumulation paradigm could be driven by either the absolute number of landmarks or the spatiotemporal density of visual stimuli. To dissociate these possibilities, we employed a Distributed Landmark paradigm in which additional landmarks were inserted while maintaining a constant spatial separation along the virtual track, thereby increasing the number of landmarks without increasing visual ensemble density (Figure 4A). Under these conditions, CA1 place-cell recruitment increased linearly (among conditions: χ²_(4)_ = 53.90, p < 0.0001; 2 vs. 4: χ²(1) = 7.35, p = 0.007; 8 vs. 10: χ²(1) = 4.3, p = 0.04; χ² tests, Bonferroni-corrected α = 0.0125), scaling continuously as more track segments became visually anchored (Figures 4B and 4C). Conversely, CA3 place-cell recruitment remained at a uniformly low baseline without the abrupt change observed in the accumulation paradigm (among conditions: χ²_(4)_ = 4.51, p = 0.25), paralleling the persistently low visual complexity of the dispersed landmarks (Figure S6A).

**Figure 4.**
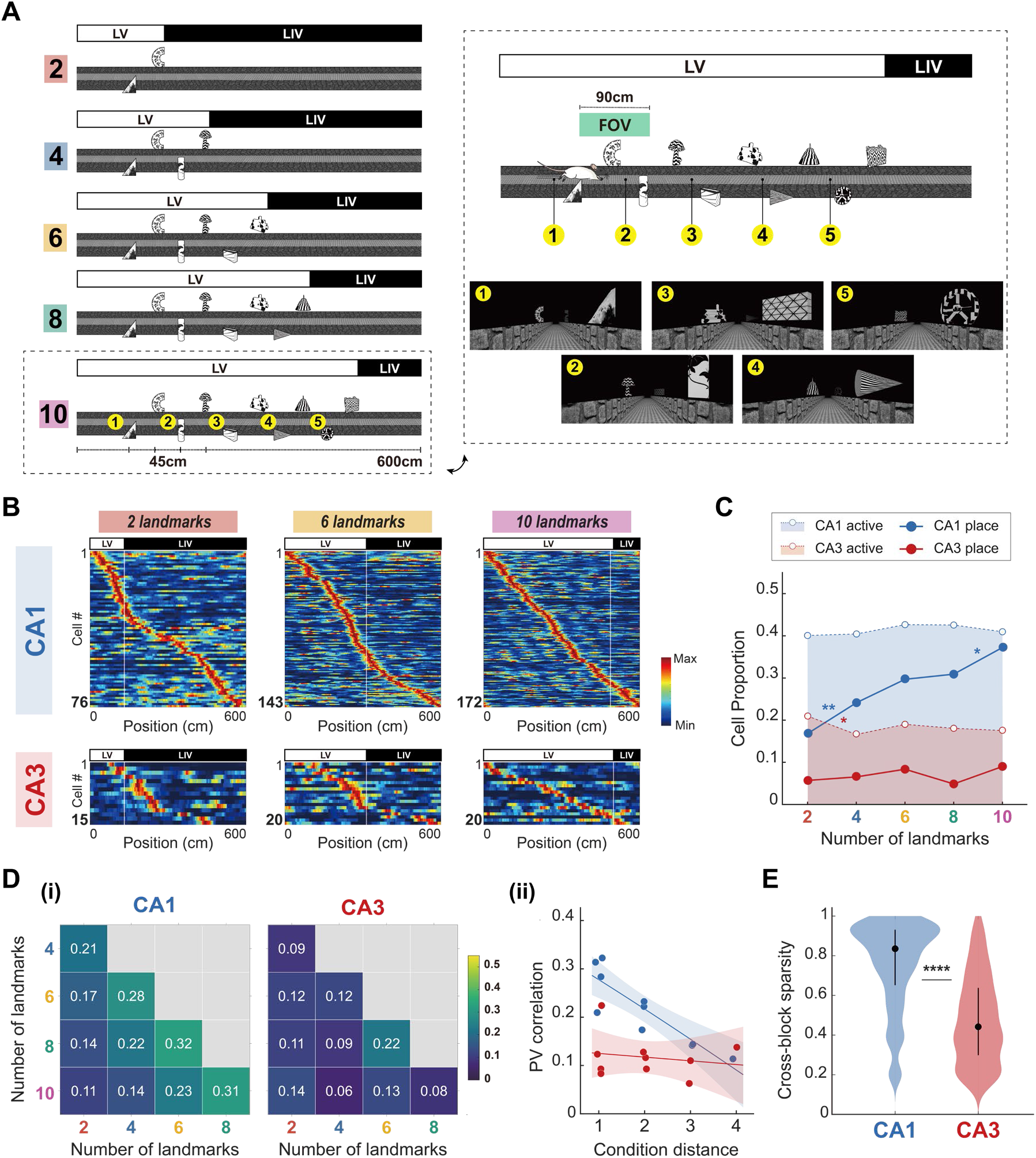
Distributed landmarks fail to induce CA3 place cell recruitment despite increasing the number of landmarks. (A) Schematic of the ‘Distributed Landmarks’ paradigm. Up to 10 landmarks were spatially dispersed across an extended 600-cm track at 45-cm intervals. Given the 90-cm FOV restriction, this configuration ensured that a maximum of three landmarks were visible at any given position, preventing the formation of a coherent visual ensemble regardless of the number of landmarks. (B) Population firing rate heatmaps for CA1 (top) and CA3 (bottom) across 5 landmark conditions. Rows represent individual place cells sorted by their peak firing location, displayed as in Figure 3A; note the markedly different cell numbers between regions. CA1 place cells more frequently formed multiple firing fields in this paradigm, in line with its extended 600- cm track. (C) Proportion of active cells (open circles with dotted line) and place cells (filled circles with solid line) in CA1 (n = 1125 units) and CA3 (n = 1270 units). The proportion of active cells remained stable across conditions in both regions (CA1: χ²(4) = 2.61, p = 0.62; CA3: χ²(4) = 8.91, p = 0.06; χ² tests). (D) PV correlation from all place cells. (i) PV correlation matrices for CA1 (left) and CA3 (right), displayed as in Figure 3D. (ii) PV correlation of every condition pair (n =10 per region) against condition distance, with linear regression on Fisher-Z transformed coefficients and 95% confidence intervals. (E) Firing sparsity of CA1 and CA3 place cells across the five blocks in the DL paradigm. Similar to the LA paradigm, CA3 exhibited significantly lower sparsity values than CA1 (CA1 vs. CA3: Z = 22.65, p < 0.0001; Wilcoxon rank-sum test), reflecting their intrinsic regional characteristics. * p < 0.05, ** p < 0.01, **** p < 0.0001

PV correlations further highlighted this dissociation (Figure 4D). The CA3 regression slope was statistically indistinguishable from zero (slope = -0.008, t_(8)_ = -0.52, p = 0.61; linear regression on Fisher-Z transformed coefficients), confirming a failure to form a continuous spatial representation in the absence of a visual ensemble while maintaining firing sparsity (Figure 4E). CA1 retained a significant distance-dependent decay (slope = -0.06, t_(8)_ = -5.19, p < 0.0001), and correlations were consistently lower in CA3 than in CA1 (difference = -0.22, t_(16)_ = -5.0, p < 0.0001; GLM main effect of Region), with a significantly steeper decay slope in CA1 (t_(16)_= 2.94, p = 0.009; GLM Region × Distance interaction). The overall correlation magnitude in CA1 was nonetheless lower than in the accumulation paradigm (difference = -0.16, t_(17)_ = -8.26, p < 0.0001; GLM main effect of Paradigm), reflecting the continuous recruitment of new CA1 place cells for each additionally anchored track segment, supported by the population center of mass dynamically tracking the direction of landmark presentation (Figure S6B). Together, these findings demonstrate that increasing landmark count alone is insufficient; CA3 activation requires spatiotemporal integration of landmarks into a coherent visual ensemble.

### Spatial configuration controls CA3 representational content

Having established that a visual ensemble is necessary for CA3 recruitment, we next asked whether CA3 merely detects the presence of an ensemble or encodes its specific spatial configuration. If CA3 is only sensitive to aggregate visual density, its map should remain stable as long as landmark identities are preserved; conversely, if it encodes relational configurations, shuffling landmark positions should drive global remapping. To test this, we implemented a Landmark Swap paradigm in which landmark positions were randomly shuffled while preserving their identities and overall density (Figure 5A and S7A).

**Figure 5.**
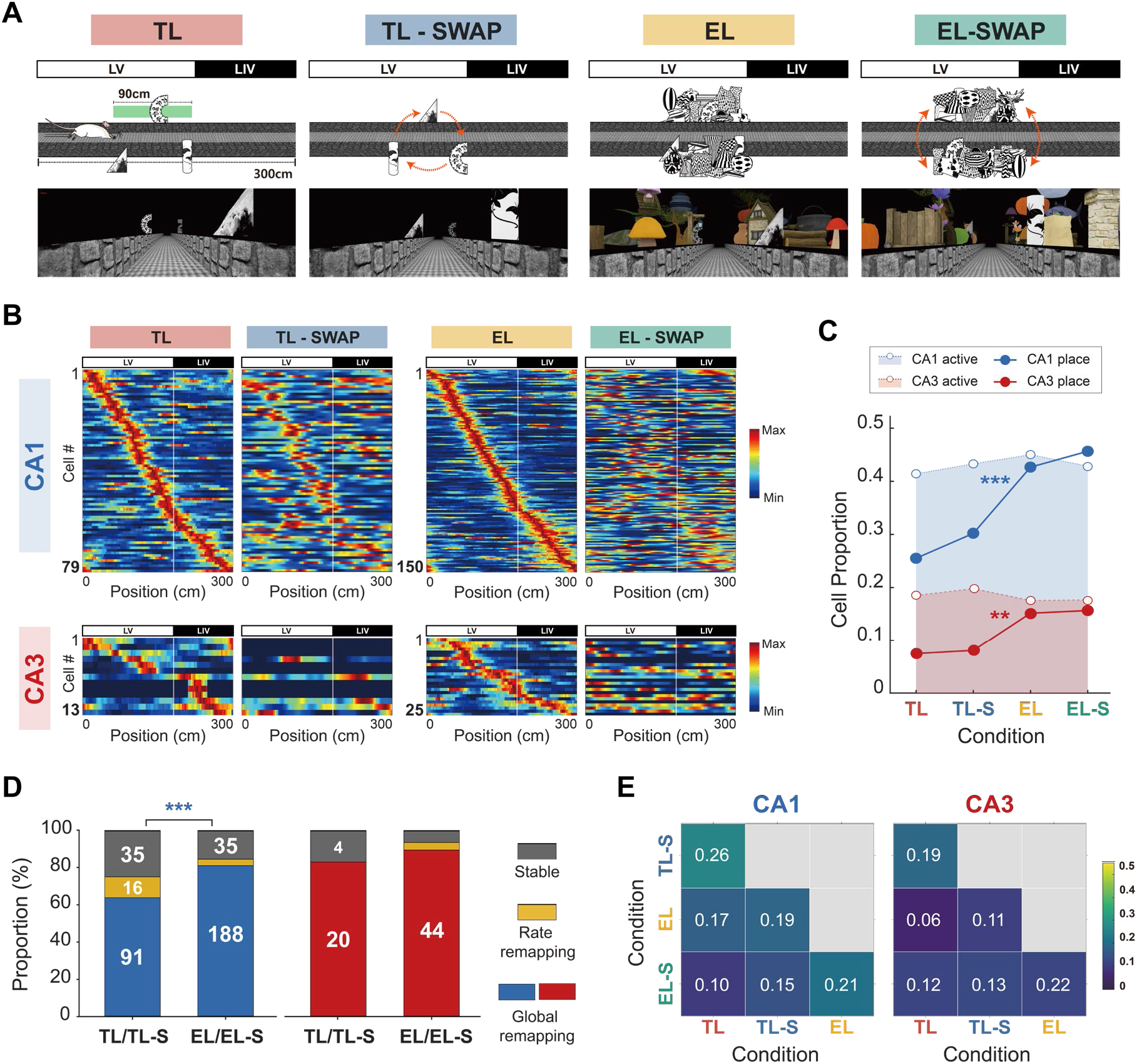
Spatial configuration of landmarks determines CA3 place cell recruitment content. (A) Schematic of the ‘Landmark Swap’ paradigm. Two landmark densities were tested: Triple Landmarks (TL) and Ensemble Landmarks (EL). In ‘Swap’ sessions (TL-Swap, EL-Swap), individual landmark identities were preserved, but their spatial positions within the LV zone were randomly shuffled. (B) Representative population firing rate heatmaps for CA1 (top) and CA3 (bottom) comparing original and swap configurations. Rows represent individual cells sorted by peak firing location in the original condition, displayed as in Figure 3A; note the markedly different cell numbers between regions. (C) Proportion of active cells (open circles with dotted line) and place cells (filled circles with solid line) in CA1 (n = 792 units) and CA3 (n = 940 units). The proportion of active cells did not differ across the four conditions in either subregion (CA1: χ²(3) = 1.41, p = 0.70; CA3: χ²(3) = 2.01, p = 0.57; χ² tests). Place cell proportions were unchanged between EL and EL-Swap configurations (CA1: χ²(1) = 0.44, p = 0.51; CA3: χ²(1) = 0.02, p > 0.99). (D) Single-unit remapping classifications (Global, Rate, and Stable) for CA1 (left) and CA3 (right) between the original and swap configurations. Cells identified as place cells in at least one of the paired conditions were included (CA1: n = 142 for TL, 231 for EL; CA3: n = 24 for TL, 49 for EL). (E) PV correlations for CA1 and CA3 across the four conditions. (i) PV correlation matrices for CA1 (left) and CA3 (right), displayed as in Figure 3D. High correlation values indicate stable spatial representations, while low values between original and swap conditions (especially in EL) indicate global remapping or orthogonalization triggered by change in spatial configuration. Significance of the difference in representational similarity between the TL and EL densities was assessed via bootstrap tests. * p < 0.05, ** p < 0.01, *** p < 0.001

Representative population maps revealed that this spatial rearrangement reorganized place-cell activity in both subregions despite constant visual complexity (Figure 5B). Place-cell proportions were significantly higher in the ensemble than in the sparse conditions for both CA1 (χ²_(1)_ = 41.79, p < 0.0001; χ² test) and CA3 (χ²_(1)_ = 11.67, p = 0.0006), yet remained unchanged between the original and swap configurations within each density (Figure 5C). The spatial content of the map, however, was profoundly reorganized (Figure 5D): CA3 exhibited predominant global remapping across both landmark densities (χ²(2) = 2.92, p = 0.23), whereas CA1 retained more stable and rate-remapping cells during the Triple Landmark (TL)-Swap than the Ensemble Landmark (EL)-Swap (χ²(2) = 16.07, p = 0.0003). Thus, the scale of network recruitment was dissociable from the content it carried.

These effects were mirrored at the population level (Figure 5E). In CA3, similarity between the original and swap configurations remained uniformly low for both TL (r = 0.19) and EL (r = 0.12) conditions (p = 0.10; bootstrap test for difference), confirming robust reorganization regardless of visual density. CA1 instead preserved higher similarity following the TL-Swap than the EL-Swap (r = 0.26 vs. 0.21, p = 0.046; Figure S7B), reflecting a reliance on individual landmarks under sparse conditions that diminished in the ensemble condition. Thus, CA3 constructs orthogonalized spatial codes with a configural selectivity tuned to the specific landmark arrangement, potentially driving the deeper reconfiguration of CA1 maps under ensemble conditions.

### Naturalistic backgrounds facilitate early CA3 recruitment

Unlike visually impoverished environments where a landmark ensemble is strictly required, natural settings feature continuous backgrounds that bind discrete objects into spatially integrated visual scenes. We hypothesized that by providing this global contextual structure, backgrounds act as a ‘spatial scaffold’ to facilitate CA3 recruitment even under sparse landmark conditions. To test this, we examined a Simple Natural Scene (SNS) environment, incrementally accumulating naturalistic landmarks against a simple, continuous background following the accumulation protocol (Figure 6A and S8A).

**Figure 6.**
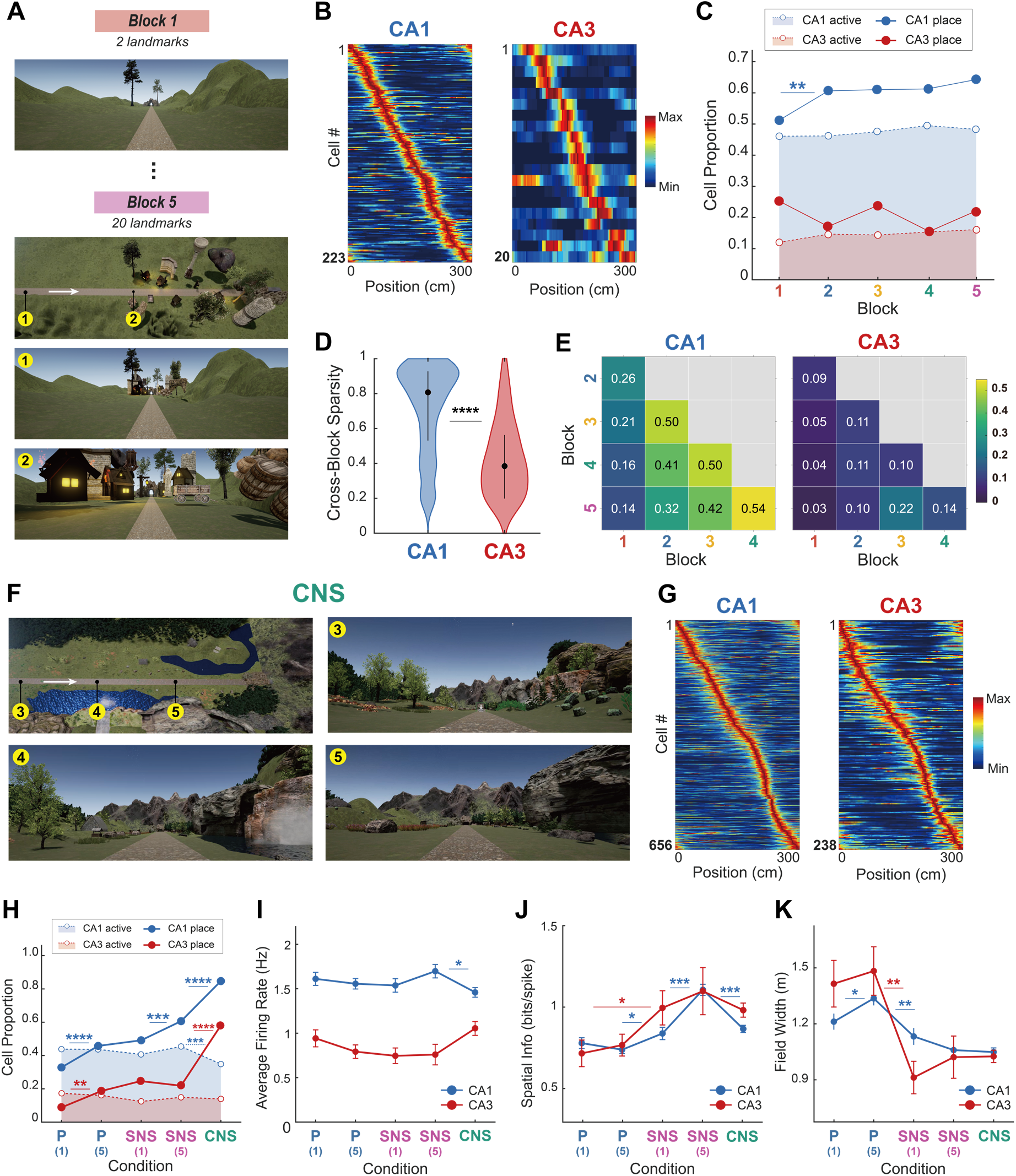
Background visual richness facilitates early and stepwise CA3 recruitment. (A) Simple Natural Scene (SNS). Schematic of the SNS environment containing naturalistic landmarks against a simple background comprising a horizon and rolling hills. The number of landmarks increased in steps of two following the accumulation protocol. (B) Population firing rate heatmaps for CA1 (left) and CA3 (right) in the SNS environment, sorted by peak firing location in Block 5, confirming consistent spatial firing along the track, displayed as in Figure 3A. (C) Proportion of active cells (open circles with dotted line) and place cells (filled circles with solid line) in CA1 (n = 870 units) and CA3 (n = 684 units) across 5 blocks in the SNS environment. (D) Comparison of Cross-Block Sparsity between CA1 and CA3 in the SNS environment. Significantly lower sparsity in CA3 indicates block-specific representations despite the background scaffold. (E) PV correlation matrices for all CA1 (left) and CA3 (right) place cells in the SNS environment, displayed as in Figure 3D. (F) Complex Natural Scene (CNS). Schematic of the CNS environment containing naturalistic landmarks against a complex, high-contrast background. (G) Population firing rate heatmaps for CA1 (left) and CA3 (right) in the CNS environment, sorted by peak firing location, displayed as in Figure 3A. (H-K) Comparisons of place field properties across the five stages of scene complexity: P-1, P-5, SNS-1, SNS-5, and the CNS, denoted P(1), P(5), SNS(1), SNS(5) on the axes (P, Poor; SNS, Simple Natural Scene; numerals denote block). (H) Proportion of active and place cells. Note the stepwise progression in CA3 versus the linear scaling in CA1. The overall fraction of active cells in CA1 decreased slightly in the CNS relative to the Poor and SNS environments (SNS-5 vs. CNS: χ²(1) = 13.4, p = 0.0003; χ² test). (I) Average firing rates (Hz) of place cells (CA1: SNS-5 vs. CNS: Z = 2.19, p = 0.03; Wilcoxon rank-sum test). (J) Spatial information scores (bits/spike) of place cells. Both subregions showed improved spatial coding across environments (CA1: χ²(4) = 81.33, p < 0.0001; Kruskal–Wallis test). CA1 exhibited progressive increases across the SNS conditions (SNS-1 vs. SNS-5: Z = -5.54, p < 0.0001; Wilcoxon rank-sum test) before a reduction in the CNS (SNS-5 vs. CNS: Z = 6.73, p < 0.0001). (K) Place field width (m). Widths narrowed significantly across the environmental gradient (CA1: χ²(4) = 48.35, p < 0.0001; CA3: χ²(4) = 19.70, p = 0.0006; Kruskal–Wallis test). Note that place fields in CA1 broadened slightly during initial landmark accumulation (P-1 vs. P-5: Z = -2.11, p = 0.03; Wilcoxon rank-sum test) before sharpening significantly upon the introduction of a background. * p < 0.05, ** p < 0.01, *** p < 0.001, **** p < 0.0001

In this naturalistic setting, robust place fields emerged in both subregions from the earliest blocks (Figure 6B and S8B). Remarkably, the proportion of CA3 place cells in SNS Block 1 already matched the peak level observed in the visually impoverished environment (Poor Block 5; χ²_(1)_ = 1.57, p = 0.21; χ² test) (Figure 6C). This indicates that the background context strongly facilitates CA3 recruitment: even the sparsest landmark condition drove CA3 activation to the level that, without a background, was reached only at the highest landmark density. This elevated CA3 recruitment remained stable across subsequent SNS blocks (among conditions: χ²_(4)_ = 4.44, p = 0.35), whereas CA1 exhibited early spatial tuning with place-cell proportions increasing significantly only across early blocks (among conditions: χ²_(4)_ = 14.21, p = 0.007; Block 1 vs. 2: χ²_(1)_ = 8.43, p = 0.003; Bonferroni-corrected α = 0.0125).

Despite this facilitated recruitment, CA3 maintained block-specific, orthogonal spatial representations, mirroring the sparsity observed without a background (Z = 10.41, p < 0.0001; Wilcoxon rank-sum test) (Figure 6D). PV correlations recapitulated this selectivity (Figure 6E). While CA1 correlations decayed with block distance (slope = -0.12, t_(8)_ = -3.08, p = 0.015; linear regression on Fisher-Z transformed coefficients), CA3 correlations remained uniformly low across all transitions (slope = -0.02, t_(8)_ = -1.51, p = 0.17). Overall, correlations were significantly lower in CA3 (difference = -0.46, t_(16)_ = -4.90, p = 0.0002; GLM main effect of Region), with CA1 exhibiting a steeper decay slope (t_(16)_ = 2.27, p = 0.037; GLM Region × Distance interaction). Thus, continuous background scenes act as a spatial scaffold to facilitate CA3 recruitment, while preserving the orthogonalized encoding of distinct landmark configurations.

### Stepwise CA3 recruitment across graded scene complexity

To probe how background complexity shapes spatial map recruitment, we introduced a Complex Natural Scene (CNS) incorporating highly enriched features such as dense forests and mountains, where both subregions formed robust place fields (Figures 6F, 6G and S8C). We then compared five stages spanning a gradient of scene complexity: P-1 (Poor Block 1), P-5, SNS-1 and SNS-5, and the CNS. Across this gradient, place cell recruitment increased substantially in both CA1 (among conditions: χ²_(4)_ = 349.97, p < 0.0001) and CA3 (among conditions: χ²_(4)_ = 167.20, p < 0.0001) (Figure 6H).

Further pairwise comparisons revealed a distinct stepwise progression of CA3 recruitment. An initial recruitment phase occurred when either a landmark ensemble or a simple background induced nascent spatial representations (P-1 vs. P-5: χ²_(1)_ = 10.61, p = 0.001; P-1 vs. SNS-1: χ²_(1)_ = 15.01, p = 0.0001; χ² test; Bonferroni-corrected α = 0.0125). CA3 recruitment then plateaued across P-5 to SNS-5 (among conditions: χ²_(2)_ = 1.60, p = 0.45), maintaining stable intermediate representations. A further recruitment phase emerged only upon transition to the CNS, where the proportion of CA3 place cells reached its highest level (SNS-5 vs. CNS: χ²_(1)_ = 45.26, p < 0.0001). By contrast, CA1 recruitment scaled almost linearly, increasing significantly at each major transition (P-1 vs. P-5: χ²_(1)_ = 25.87, p < 0.0001; SNS-1 vs. SNS-5: χ²_(1)_ = 10.88, p = 0.001; SNS-5 vs. CNS: χ²_(1)_ = 82.63, p < 0.0001).

Spatial coding quality improved as background scenes were introduced (Figure S8D), while basic firing-rate properties and running speeds remained stable across environments (Figures 6I and S8E). Specifically, the presence of a background scaffold significantly increased spatial information scores compared to visually impoverished environments (CA1: P-5 vs. SNS-1: Z = -2.05, p = 0.04; CA3: P-1 vs. SNS-1: Z = -2.11, p = 0.034; Wilcoxon rank-sum test) (Figure 6J). Spatial tuning also sharpened, as evidenced by significantly narrower place-field widths upon transitioning to the naturalistic environment in both subregions (P-5 vs. SNS-1; CA1: Z = 2.98, p = 0.003; CA3: Z = 2.61, p = 0.009) (Figure 6K). These results demonstrate that background scaffolds not only facilitate the initial recruitment of hippocampal maps but also refine the spatial precision of their representations.

### CA3 recruitment shifts fast- to slow-gamma coupling

While single-unit analyses revealed that CA3 map recruitment follows a stepwise progression—initiated by the nonlinear gating of visual ensembles and further facilitated by background scaffolds—the underlying circuit-level dynamics coordinating these transitions remain unresolved. We analyzed LFPs, focusing on theta-gamma phase-amplitude coupling (PAC) as electrophysiological readouts for external entorhinal inputs (fast gamma, FG; 65- 100 Hz) and internal CA3 processing (slow gamma, SG; 30-50 Hz) (Figure 7A)^9,35^. As landmark density reached its maximum in P-5, FG-PAC in CA3 decreased significantly (Figures 7B and 7C). This attenuation temporally coincided with the initial nonlinear state transition of the CA3 spatial map (Block 1-4 vs. Block 5: β = -0.0002, p = 0.0025; linear mixed-effects model) (Figure 7D). In contrast, SG-PAC in CA3, alongside overall PAC levels in CA1, remained relatively stable (Figure S9A, B). To further determine how scene complexity influences these network states across environments, we compared PAC magnitudes across the Poor, SNS, and CNS environments (Figures 7E–7G). While SG coupling remained at baseline levels in both the Poor and SNS environments, it exhibited a substantial increase in the CNS (P-5 vs. CNS, Z = -4.71, p < 0.001; SNS-5 vs. CNS, Z = - 3.97, p < 0.001; Wilcoxon rank-sum test) (Figure 7G)—a transition absent in CA1 (Figure S9C).

**Figure 7.**
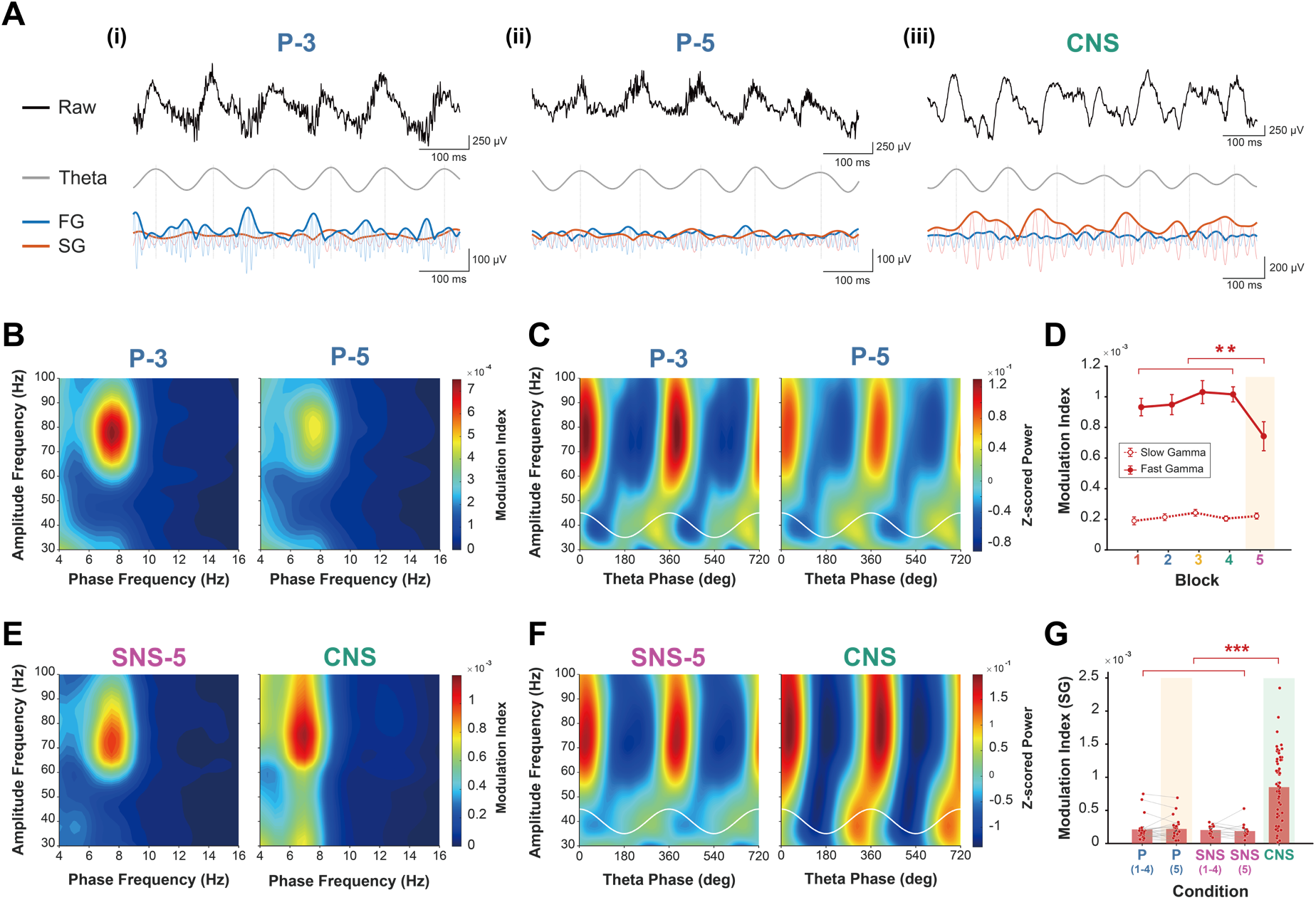
Theta-gamma phase-amplitude coupling reveals transition from fast gamma decoupling to slow gamma surge in CA3. (A) Representative CA3 LFP traces for (i) Block 3 (P-3) and (ii) Block 5 (P-5) in the Landmark Accumulation paradigm, and (iii) the CNS environment, confirming the dynamic engagement of distinct gamma bands during spatial navigation. From top to bottom: raw LFP (black), filtered theta (gray), fast gamma (blue), and slow gamma (orange). In the gamma traces, faint and bold lines indicate filtered LFP and Hilbert transform envelopes, respectively. (B) Phase-amplitude comodulograms for Blocks 3 and 5 in the Landmark Accumulation paradigm. (C) Theta-resolved gamma spectra for Blocks 3 and 5 in the Landmark Accumulation paradigm. (D) Modulation Index (MI) of slow gamma (dashed line) and fast gamma (solid line) across Blocks 1 to 5 in the Landmark Accumulation paradigm. Orange shading indicates Block 5 where the initial CA3 place cell recruitment occurs. Error bars indicate within-subject SEM. (E) Same as (B), but for Block 5 in the SNS and CNS environments. (F) Same as (C), but for Block 5 in the SNS and CNS environments. (G) MI of slow gamma across the five stages of scene complexity. Bars and error bars represent the mean and the between-subject SEM. Single dots indicate individual sessions, and gray lines connect paired sessions. Orange and green shadings indicate conditions where the stepwise activation of CA3 occurs. ** p < 0.01, *** p < 0.001

Together, these LFP dynamics reveal two distinct network-level signatures of CA3 map recruitment. The first—a selective attenuation of FG-PAC at P-5—coincides with the initial nonlinear state transition, indicating a state of sensory decoupling wherein the CA3 suppresses external EC inputs. The second—an abrupt increase in SG-PAC specifically in the CNS—accompanies the further increase in CA3 activation driven by maximal scene complexity, reflecting the enhanced recurrent processing. Although their sequential relationship was not observed within a single session, their co-alignment with the two-stage CA3 recruitment across environments (Figure 6H) supports a stepwise model: initial subnetwork activation and EC decoupling (Stage 1) precede the global network engagement enabled by rich scene context (Stage 2; see Figure S10 and Discussion).

## DISCUSSION

While remapping and place-field formation have been extensively characterized, how the hippocampus initially recruits a spatial map from a quiescent network—a distinct computational problem we term ‘recruitment’—remained largely unexamined. By parametrically manipulating visual landmark density, spatial configuration, and scene complexity in a VR environment while recording simultaneously from CA1 and CA3, we show that these subregions follow qualitatively different recruitment rules, establishing recruitment as a third, mechanistically distinct level of hippocampal spatial coding. While CA1 continuously updates with landmark accumulation, CA3 remains functionally quiescent until a spatiotemporally integrated visual ensemble triggers a nonlinear state transition, encoding the configuration as an orthogonalized spatial representation. LFP dynamics further reveal that this transition unfolds in a stepwise manner, progressing from selective sensory decoupling to global recurrent engagement as scene complexity increases. These findings provide a circuit-level account of how CA3 and CA1 divide the labor of constructing the cognitive map.

### Beyond remapping and formation: the recruitment problem

The findings presented here address the ‘recruitment problem’—the macroscopic transition of a quiescent network into a functional spatial representation^18,19,36^. Because animals in most prior studies encounter richly cued environments from the outset, observing a genuine quiescent baseline to dissociate the initial recruitment conditions of different hippocampal subregions has been difficult. By establishing these requisite conditions, we demonstrate that this subregional divergence is a population-level phenomenon distinct from independent single-cell plasticity that underlies place-field formation. Furthermore, this fundamental difference is obscured in standard remapping paradigms, where both regions respond to environmental changes from an already-active baseline^7,12^, underscoring the importance of investigating recruitment as a distinct phenomenon.

### Spatiotemporal integration and synchronous activation of CA3 subnetworks

The critical trigger for CA3 recruitment is not landmark count *per se*, but the availability of a spatiotemporally integrated visual ensemble (Figures 3 and 4)—defined as a cluster of landmarks that are simultaneously visible and temporally contiguous within the restricted FOV. Within the CA3 autoassociative architecture^11,37^, recurrent collaterals require coordinated co-activation of a neuronal population to transition into a new state^3,10^. Consequently, sparse landmarks fail to provide sufficient excitatory drive to overcome baseline local inhibition^38^. However, a visual ensemble delivering spatiotemporally integrated inputs provides the convergent excitation necessary to evoke the synchronous discharge of highly interconnected CA3 subnetworks^37^. This synchrony is evidenced by significantly increased pairwise spike correlations rather than independent single-unit bursting (Figure S5).

Because BTSP-like mechanisms in CA3 require stronger dendritic depolarization and exhibit lower induction probabilities than in CA1^16,17,39–41^, this convergent, synchronized input is likely essential to provide the sustained depolarization needed to induce such plasticity. Although we did not directly measure dendritic dynamics or synaptic plasticity here, the abrupt, synchrony-dependent nature of CA3 recruitment supports this interpretation, generating a testable prediction for future intracellular studies.

This synchronous discharge of CA3 subnetworks drives distinct microcircuit dynamics with dual network-level consequences. First, recurrent excitation within these subnetworks amplifies the synchronized depolarization, driving the nonlinear state transition. Concurrently, this synchronous activation triggers a rapid disynaptic feedback inhibitory surge^42^, creating a functional barrier against subsequent feedforward excitation. Consistent with this, the sharp decrease in FG-PAC—which conveys real-time external sensory inputs from the EC^35^—coincides precisely with ensemble-driven recruitment (Figure 7). This sensory decoupling indicates that once the initial spatial map is established, the CA3 network actively suppresses further external streams. By shifting toward an internally driven, high-inhibition state, CA3 protects the nascent spatial representation from external noise.

### Configural selectivity reveals relational encoding in CA3

The robust orthogonalization of the CA3 map following spatial rearrangement (Figure 5) indicates that CA3 evaluates the configural structure of the environment, binding independent elements into a conjunctive representation rather than merely aggregating individual landmarks^43–45^. Consequently, spatial rearrangement disengages the previously active subnetwork and recruits a distinct one. CA1 may act as a comparator that receives both CA3 and EC inputs^4,5^, exhibiting a representational shift driven by the reconfigured CA3 output.

In the SNS environment, continuous background scaffolding facilitated CA3 recruitment even with sparse landmarks (Figure 6). This is consistent with a subthreshold priming mechanism: sustained background input—plausibly conveyed via EC projections encoding the global spatial context^46–48^—elevates baseline excitability in CA3 dendrites, lowering the threshold for ensemble-driven subnetwork activation. Within this primed state, sparse landmarks can reliably trigger recruitment. Because the background provides a spatially non-specific drive, spiking output remains gated by the specific configuration of local landmarks, preserving high representational sparsity. Although we did not directly measure EC activity or dendritic potentials, this interpretation aligns with the broad temporal integration window of CA3 recurrent circuits^49,50^ and generates a clear prediction: EC inactivation should abolish this background-mediated scaffolding effect.

Upon transitioning to the CNS environment, convergent excitatory drive likely reaches a critical mass of the recurrent network^3,10^. Exceeding this activation condition shifts the excitation-inhibition balance toward sustained recurrent excitation, triggering global network engagement as reflected by the SG-PAC increase (Figure 7). Thus, CA3 map recruitment follows a stepwise trajectory: from initial subnetwork activation and sensory decoupling to global network engagement and recurrent synchronization.

We rule out two alternative explanations. First, the abrupt CA3 recruitment does not merely reflect a novelty effect, because the decreasing sequence (20-to-2 landmarks) yielded an indistinguishable recruitment profile anchored to ensemble density rather than the initial novel condition (Figure S4D). Were novelty the driver, the first condition of the decreasing sequence—the earliest encountered—should itself show elevated CA3 recruitment, which it did not. Furthermore, early CA3 recruitment in the SNS environment occurred before any landmark manipulation, ruling out a novelty explanation for the scaffolding effect. Second, CA3 quiescence in sparse conditions is not an artifact of recording instability, as units silent during navigation fired reliably during surrounding sleep periods (Figure S2). Moreover, their orthogonal encoding of specific configural arrangements (Figure 5) demonstrates a genuine, configuration-sensitive computational threshold rather than a non-specific response to increased visual stimulation.

### Background scaffolding and the circuit-level basis of scene construction

The progressive facilitation of CA3 recruitment across the Poor → SNS → CNS hierarchy provides direct rodent neurophysiological evidence bearing on SCT^27,51^. Our results operationalize this construct: for the CA3 autoassociative network, a ‘scene’ can be defined as a configuration of landmarks embedded within a coherent spatial background that together provide sufficient convergent excitation to cross the threshold for attractor state formation. A mere accumulation of independent objects, lacking relational and contextual structure, does not suffice.

This circuit-level formulation provides a specific neural mechanism—threshold-gated CA3 attractor recruitment—for the qualitative difference between object-based and scene-based processing observed in human imaging^25,32,33^ and rodent behavior^28–31^. Furthermore, by attributing the scene construction function specifically to CA3 while assigning a distinct landmark-integration role to CA1, our results establish a subregional division of labor unaddressed by original formulations of SCT.

Although our paradigm did not parametrically isolate scene complexity as a single independent variable—conflating changes in background richness, naturalistic landmark character, and overall visual entropy—future work systematically grading these individual components will be necessary to disentangle these variables and further refine the operational definition of a ‘scene’ in terms of hippocampal circuit dynamics.

### Complementary relationships between CA3 and CA1: Balancing orthogonalization with generalization

The differential recruitment dynamics reported here support a complementary model for constructing and maintaining the cognitive map (Figure S10)^52–54^. CA3 operates as a nonlinearly gated ‘scene constructor’. By remaining quiescent until a coherent structural context emerges and then decoupling from external noise, it prevents unstable mapping and yields distinct, orthogonalized spatial representations. Conversely, CA1 functions as a ‘continuous integrator’ that progressively accumulates landmark information. It maintains generalized spatial tuning to provide an uninterrupted, provisional spatial reference in impoverished environments, supporting flexible navigation^55^.

This subregional synergy is mediated by the CA1 comparator mechanism^4,5,56–58^. By receiving direct inputs from the EC, CA1 evaluates ongoing sensory evidence against internal contextual frameworks provided by CA3^46,59^. Prior to CA3 recruitment, the provisional map in CA1 likely corresponds to the fragmented representations observed during early environmental exploration^19,60^. Upon ensemble-driven CA3 recruitment, the resulting attractor output stabilizes and reorganizes these CA1 representations^61^. Furthermore, the inverted U-shaped patterns across the SNS-to-CNS transition reflect an essential functional trade-off: highly enriched contexts facilitate global map stability and spatial coverage at the expense of maximal single-unit specificity (Figure 6J). This optimization is likely essential for maintaining a reliable cognitive map in highly complex, real-world scenes.

This proposed framework leaves several questions for future investigation. First, the causal role of CA3 attractor engagement in stabilizing CA1 representations requires direct testing via pathway-selective interventions. Second, the specific upstream contributions of the lateral and medial entorhinal cortices^47,48,62–64^ in driving CA3 subnetwork activation must be dissected. Third, it remains unknown whether repeated exposure to visually impoverished environments gradually facilitates CA3 recruitment through experience-dependent plasticity, or if the requisite conditions remain fixed. Finally, as our body-restrained VR paradigm isolates visually driven map formation, future studies must determine how self-motion cues and multisensory convergence interact with visual ensembles to govern CA3 recruitment in freely moving animals.

## MATERIALS AND METHODS

### Subjects

A total of 20 male Long–Evans rats (400–500 g, 18–24 weeks old) were used. Eight rats were newly recorded for the present experiments, and data from the remaining 12 were obtained from a previously published study^34^ to enable direct comparison with the Complex Natural Scene (CNS) environment. Animals were individually housed under temperature- and humidity-controlled conditions with a 12-h light/dark cycle (lights on at 8:00 AM). Rats had *ad libitum* access to food, whereas water intake was restricted to maintain body weight at ∼85% of the free-feeding level. All procedures were approved by the Institutional Animal Care and Use Committee of Seoul National University (SNU-230427-1-6).

### Hyperdrive implantation surgery

Each rat was implanted with a custom-designed hyperdrive containing a dual-bundle arrangement of 24 tetrodes and three reference electrodes, configured for maximal coverage along the proximodistal axis of the dorsal hippocampus. Rats were initially anesthetized with an intraperitoneal injection of sodium pentobarbital (Nembutal, 65 mg/kg), and anesthesia was maintained with isoflurane (0.5–2% in 100% oxygen) throughout the surgery. The hyperdrive was positioned over the right hippocampus, centered 3.8 mm posterior to bregma and 3.0 mm lateral to the midline. After at least one week of recovery, tetrodes constructed from 17-μm platinum–iridium wires were gradually advanced over several days to reach the targeted hippocampal subregions.

### Electrophysiological recording

Neural signals were recorded with a Digital Lynx SX acquisition system (Neuralynx) via a headstage (HS-36, Neuralynx) connected to an electrode interface board (EIB-36-24TT, Neuralynx) mounted on the hyperdrive. Signals were band-pass filtered (0.6–6 kHz), sampled at 32 kHz, amplified 1,000–10,000-fold, and thresholded to detect spikes. Recordings targeted the pyramidal layers of dorsal CA1 and CA3. To achieve simultaneous coverage, individual tetrodes were assigned to CA1 or CA3 based on their mediolateral and proximodistal positions within the bundle. Tetrode depths were finely adjusted using physiological markers, such as sharp-wave ripples during sleep, according to established criteria.

### VR system and experimental apparatus

The virtual reality (VR) environment was rendered in Unreal Engine 4.14 (Epic Games) and displayed on five 24-inch LCD monitors, providing a ∼300° horizontal and -45° to +75° vertical field of view (FOV). Rats were body-restrained in a custom-made jacket and positioned on a Styrofoam cylinder (40 cm diameter, 20 cm width) covered with a commercial yoga mat to ensure adequate friction. Cylinder rotation was measured by a high-resolution rotary encoder (DBS60EBGFJD1024; Sick) mounted on the supporting frame and converted to virtual displacement via a custom MATLAB (MathWorks)–Arduino interface. Water rewards (20 μl) were delivered through a motorized lick port (L16-R; Actuonix Motion Devices) that extended toward the snout only during reward delivery, preventing its use as a stable spatial cue. Licks were detected by an infrared sensor (FD-S32; Panasonic), which triggered a solenoid valve (VA212-3N; Aonetech) to dispense water. Reward timing and lick- port motion were controlled by Unreal Engine via the Arduino interface.

To synchronize behavioral and electrophysiological data, an Arduino-generated transistor– transistor logic (TTL) pulse was emitted for each rendered VR frame (30 Hz) and recorded by the Neuralynx system, enabling frame-accurate alignment of VR and neural timestamps.

### Training schedule

#### (i) Pre-surgical training and familiarization

Prior to surgery, rats underwent pre-training to acclimate to the VR apparatus and 1D track navigation. Familiarization sessions were conducted in a 300-cm virtual linear track featuring a grayscale 2D scene (the harbor at San Diego Bay). Rats learned to traverse the track to receive water rewards, which were delivered at random locations to prevent the use of reward-predictive cues as spatial anchors. Rats were trained daily for 30 min. Only animals that achieved a stable running distance of at least 360 m (120 trials) in a 30-min session on two consecutive days were advanced to the recording phase.

#### (ii) Post-surgical training and recording schedule

Following hyperdrive implantation, rats were allowed at least one week of recovery with a*d libitum* access to food and water. Water restriction was then reinstated, and post-surgical training resumed in the familiarization environment to re-establish stable running behavior and verify that the hyperdrive did not impede movement. Once pre-surgical performance levels were recovered, main recording sessions commenced. The newly implanted rats were recorded in the Poor and SNS environments in that order, whereas the previously published cohort performed the CNS followed by the Poor environment. In the Poor environment, rats sequentially performed the Landmark Accumulation, Landmark Swap, and Distributed Landmark paradigms. For all parametric sessions, the increasing sequence (e.g., 2-to-20 landmarks) was always conducted first, with the corresponding decreasing sequence (e.g., 20- to-2) tested on the subsequent recording day.

### VR environments

#### (i) Visually Poor environment

The visually Poor environment served as the baseline condition for examining CA3 recruitment across paradigms (Figures 1, 4, 5). The virtual track (300 cm for accumulation and swap paradigms; 600 cm for the distributed paradigm) was embedded in a void (black) background devoid of distal cues. A repetitive, low-contrast brick pattern on the side walls provided optic flow. Landmarks consisted of artificial 3D objects (for example, striped pillars, dotted triangular cones) with distinct shapes and textures to ensure high visual salience and discriminability. Water rewards were delivered at random locations along the track. Upon reaching the track end, rats were instantaneously teleported back to the start to minimize the use of end-of-track cues.

#### (ii) Parametric Landmark Accumulation paradigm

In the accumulation paradigm, visual landmarks were confined to a 90-cm segment within the LV zone, creating a spatiotemporally controlled window of visual input. Within each session, the number of landmarks in this segment was adjusted every 15 trials (total 150 trials). In the increasing session, the number of landmarks ramped from 2 to 20; in the decreasing session, the sequence was reversed. Condition transitions were seamless, and additional landmarks appeared at previously unoccupied coordinates. This design ensured that changes in hippocampal activity were attributable specifically to changes in landmark density rather than the introduction of a novel environment.

#### (iii) Distributed Landmark paradigm

To test whether a spatiotemporal visual ensemble is required for CA3 activation, we designed a Distributed Landmark paradigm using a 600-cm track. The total landmark count was parametrically varied across five conditions (2, 4, 6, 8, and 10 landmarks), presented in either increasing or decreasing sequence. Landmarks were placed at a constant 45-cm spacing along the track. In combination with the 90-cm window of FOV restriction, this arrangement ensured that no more than three landmarks were visible simultaneously, thereby maintaining sparse visual input and preventing ensemble formation even at the highest landmark counts.

#### (iv) Landmark Swap paradigm

To assess the contribution of spatial configuration to CA3 recruitment, we implemented a Swap paradigm. Along the 300-cm track, both the landmark segment and FOV were fixed at a 90-cm window, as in the accumulation paradigm. Each session comprised four sequential conditions: Triple Landmark (TL), TL-Swap, Ensemble Landmark (EL), and EL-Swap, using the same landmarks and configurations as in Shin et al.^34^. In Swap conditions (TL-Swap and EL-Swap), landmark identities were preserved but their relative positions were randomly shuffled within the LV segment.

#### (v) Simple Natural Scene (SNS) environment

To probe how background scenes influence CA3 recruitment, we designed a Simple Natural Scene (SNS) environment and recorded from a subset of newly implanted rats (n = 5). Here, naturalistic landmarks (for example, trees and rocks) were presented against a simple distal background of undulating hills. Unlike the Poor environment, no FOV restriction was applied to the distal background; rats could continuously view the horizon along the track. The number of landmarks was increased or decreased in steps of 2, following the same parametric accumulation protocol as in the Poor environment (Figure 1). Landmarks were confined to a 120-cm segment in the latter portion of the 300-cm track. Optic flow was provided by the texture of the hills and ground.

#### (vi) Complex Natural Scene (CNS) environment

To examine the impact of scene complexity, we used the Complex Natural Scene (CNS), a highly enriched forest environment containing numerous naturalistic landmarks embedded in a complex, high-contrast background, as in Shin et al.^34^. For this condition, we analyzed data from 12 previously recorded rats. Cell-proportion analyses were restricted to the subset of these animals that had also been recorded in the Poor environment, where they exhibited the same functionally quiescent CA3 activity. The stable, high-density CA3 recruitment exhibited by these animals in the CNS confirmed that the sparse activation observed in the Poor environment reflected the visually impoverished context rather than intrinsic physiological limitations.

### Histological procedures

After completion of recordings, final tetrode positions were marked by passing a small electrolytic current (10 μA for 10 s) through each tetrode. Rats were euthanized with CO₂ overdose and transcardially perfused, first with phosphate-buffered saline (PBS) delivered by syringe and then with 4% formaldehyde using a peristaltic pump (Masterflex Easy-Load II; Cole-Parmer). Brains were extracted and post-fixed in 4% formaldehyde with 30% sucrose at 4 °C until they sank. Coronal sections (40 μm) were cut on a freezing microtome (HM 430; Thermo Fisher Scientific), mounted on subbed slides, and Nissl-stained with thionin. High-resolution images were acquired with a fluorescence microscope (Eclipse 80i; Nikon) and a slide-scanning system (MoticEasyScan; Motic). Tetrode tracks and recording sites were reconstructed by identifying electrolytic lesions and comparing serial sections with the original hyperdrive configuration and depth profiles recorded during the sessions.

### Construction of flat map

To determine the precise proximodistal positions of recording sites (Figure S2), we constructed a linearized flat map of the dorsal hippocampal pyramidal layer following established methods. High-resolution images of serial Nissl-stained coronal sections were aligned in Fiji (ImageJ), and the curvilinear trajectory of the pyramidal cell layer was reconstructed by marking closely spaced points along the layer in each section. To account for inter-animal variability in brain size and cutting angle, traced distances and relative positions were normalized to a standard coordinate system based on the rat brain atlas of Paxinos and Watson (2009)^65^. The proximal boundary at the dentate gyrus–CA3c junction was defined as position 0, and the distal end of the CA1 pyramidal layer as position 1^66–68^. Subfield boundaries (CA3/CA2, CA2/CA1) were identified using cytoarchitectonic criteria, including cell-body size, laminar morphology, and mossy-fiber distribution^2,66,69,70^. Final tetrode locations, identified by electrolytic lesions, were projected onto this standardized 0–1 scale.

### Unit isolation

Single units were isolated using custom spike-sorting software (WinClust) following previously described procedures^34^. Clusters were manually defined based on waveform features, including peak amplitude, valley, and energy, across the four channels of each tetrode. Cluster quality was verified by clear separation in multidimensional feature space and by the presence of a refractory period >1 ms in inter-spike interval histograms. Only units exhibiting stable waveforms and consistent firing across pre-sleep, navigation, and post-sleep epochs were included in subsequent analyses (Figure S2).

### Single-unit analysis

#### (i) Characterizing basic firing properties

For each unit, mean firing rate was computed as total spikes divided by session duration. To restrict analyses to active navigation, only spikes emitted when running speed exceeded 5 cm/s, measured by rotary encoders, were included. The peak firing rate was defined as the maximum value across spatial bins in the raw rate map. Units with mean firing rate >0.3 Hz and peak firing rate >1 Hz in at least one condition were classified as active and considered for place-cell analysis.

#### (ii) Rate map construction

One-dimensional occupancy-normalized firing-rate maps were constructed for each unit. The virtual track (300 or 600 cm) was divided into 3-cm bins. The firing rate in each bin was calculated as the spike count divided by occupancy time. Raw rate maps were smoothed using an adaptive binning procedure^71^.

#### (iii) Cross-block sparsity index

To quantify representational specificity across landmark blocks, we computed a cross-block sparsity index. For each unit, sparsity was defined as:

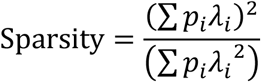

where *i* indexes blocks, *p_i_* is the occupancy probability in block*_i_*, and *λ_i_* is the mean firing rate in block*_i_*. Values near 1 indicate relatively uniform firing across blocks, whereas lower values indicate activity concentrated in a subset of blocks, reflecting higher specificity or orthogonality.

#### (iv) Quantification of spatial properties

Spatial selectivity was quantified using the spatial information (SI) score^71^:

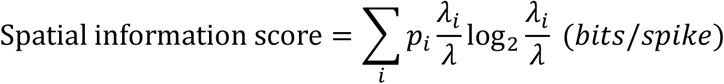

where *i* indexes bins, *p_i_* is the occupancy probability in bin*_i_*, *λ_i_* is the mean firing rate in bin*_i_*, and *λ* is the overall mean firing rate. The statistical significance of SI score was assessed with 1,000 circular-shift shuffles of the spike train. A unit was classified as a spatially informative cell (place cell) if its SI was ≥ 0.15 bits/spike and exceeded the 95th percentile of its respective shuffled distribution.

#### (v) Alluvial diagram and retrospective state classification

To track functional state transitions, each cell in each block was categorized as (1) place cell, (2) active non-spatial cell, or (3) silent cell. Alluvial diagrams were used to visualize the retrospective transitions of units identified as place cells in Block 5 (the reference block) back to their states in preceding blocks. This allowed for the quantification of the final spatial map that emerged *de novo* versus through refinement of pre-existing representations.

#### (vi) PV correlation analysis

To assess population-level representational similarity across blocks, we performed population vector (PV) correlation analysis. For each spatial bin_j_, a population vector was constructed, with elements corresponding to the mean firing rates of individual units in that bin. The PV correlation for a given bin was defined as the Pearson correlation coefficient between vectors from two blocks (A and B). Overall similarity was obtained by averaging correlations across bins. For statistical comparisons (for example, remapping or decay), correlation coefficients underwent Fisher z-transformation. Only units classified as place cells in at least one of the compared blocks were included in PV calculations.

#### (vii) Detection of place-field boundaries

Place fields were defined as contiguous spatial bins in which firing rate exceeded 20% of the peak rate for at least five consecutive bins. Field boundaries were anchored at positions where the peak firing rate within the field exceeded 1 Hz. For units with multiple fields, the field with the highest peak rate was designated as the primary place field.

### Local field potential analysis

#### (i) Preprocessing

Raw local field potential (LFP) signals were downsampled from 32 kHz to 2 kHz (Rate Reducer; Neuralynx) for all phase and amplitude analyses. To select a representative channel for each subregion (CA1 and CA3), the power spectral density was computed using a multi- taper method (Chronux Toolbox, MATLAB). The tetrode exhibiting the highest average theta power, excluding those with electrical artifacts, was selected as the reference channel. Specific frequency bands—theta (6-12 Hz), slow gamma (SG, 30-50 Hz), and fast gamma (FG, 65-100 Hz)—were extracted using a zero-phase third-order Butterworth band-pass filter (*filtfilt*, MATLAB) to prevent phase distortion. To isolate neural activity during active navigation and prevent movement-related edge artifacts, subsequent LFP analyses were restricted to periods where the animal’s running speed exceeded 5 cm/s. Continuous data were filtered prior to applying the velocity mask to ensure the temporal integrity of the filtered signals.

#### (ii) Phase-amplitude coupling (PAC)

To quantify the intensity of cross-frequency coupling between the theta phase and gamma amplitude (SG and FG), we computed the Modulation Index (MI) as described by Tort et al.^72^. Briefly, the instantaneous phases and amplitude envelopes of the filtered LFP signals were extracted using the Hilbert transform. The continuous theta phase was divided into 18 bins (20° each), and the mean gamma amplitude within each phase bin was calculated and normalized to create a phase-amplitude distribution. The MI was then defined as the Kullback-Leibler (KL) distance between this empirical distribution and a uniform distribution, divided by the logarithm of the number of bins. To assess the statistical significance of the coupling, a surrogate Z-scored MI was computed. A null distribution of MI values was generated by introducing random temporal shifts between the phase and amplitude time series, and the raw MI was Z-scored against this surrogate distribution.

#### (iii) Phase-amplitude comodulogram

Phase-amplitude comodulograms were constructed to dynamically visualize PAC across a continuous frequency space. MI values were systematically computed across a fine two-dimensional grid of phase frequencies (4-16 Hz, 0.5-Hz steps) and amplitude frequencies (30- 100 Hz, 2-Hz steps). The resulting 2D matrix was smoothed using a 2D gaussian filter to enhance visualization of the coupled frequency bands.

#### (iv) Theta-resolved gamma power spectrum

To visualize phase-locked gamma bursts while controlling for 1/f power drop-off, a time-domain Z-scoring approach was adopted^35^. The LFP signal was narrow-band filtered across amplitude frequencies (20-110 Hz, 2-Hz steps). For each frequency, the instantaneous power (squared Hilbert amplitude) was extracted and Z-scored across the entire valid epoch in the time domain. These time-domain Z-scored power values were then averaged within 36 concurrent theta phase bins (10° each), with 0° defined as the peak of the local theta cycle. For visualization, the phase axis was duplicated to display two full cycles (0-720°), and the amplitude frequency axis was constrained to 30-100 Hz.

### Quantification and statistical analysis

All analyses were performed in MATLAB 2025a (MathWorks). Chi-square tests were used to compare proportions of active cells and place cells. Wilcoxon rank-sum tests were used to compare firing rates, SI scores, cross-block sparsity, and PAC values between regions or environmental conditions. Wilcoxon signed-rank tests were used to evaluate paired within-session state transitions in LFP metrics. Right-tailed one-sample t-tests evaluated whether PAC surrogate Z-scores significantly exceeded chance level (zero). For multi-stage comparisons across environments (P-1 through CNS), Kruskal–Wallis tests were applied followed by Dunn’s *post hoc* tests with Bonferroni correction. Representational similarity decay over distance was modeled using linear regression. To compare decay slopes and overall magnitudes between regions or paradigms, an analysis of covariance (ANCOVA) was employed within a general linear model (GLM) framework, assessing interaction terms and main effects, respectively. Longitudinal trends in LFP metrics were assessed using linear mixed-effects models (LMMs) with condition block as a fixed effect and session as a random intercept. Configural sensitivity in the Landmark Swap paradigm was assessed by bootstrap resampling (N = 1,000). Unless otherwise indicated, statistical significance was set at α = 0.05, and error bars represent the standard error of the mean (SEM). To accurately isolate within-subject state transitions in LFP trajectories, within-subject SEM (Cousineau-Morey method) was used, whereas between-subject SEM was used for absolute magnitude comparisons.

## ACKNOWLEDGEMENTS

This research was supported by the National Research Foundation of Korea Grants 2019R1A2C2088799, 2022M3E5E8017723, RS-2024-00452391, RS-2025-02303740, and the Mid-Career Bridging Program through Seoul National University for Inah Lee. We thank Jhoseph Shin and Hyun-Woo Lee for data collection.

## AUTHOR CONTRIBUTIONS

E.-H.L. designed the research, collected and analyzed data, and wrote the manuscript; I.L. supervised all aspects of the research and wrote the manuscript.

