## Supplementary figures for "How visual scenes recruit spatial maps: threshold-gated activation in CA3 versus continuous integration in CA1"

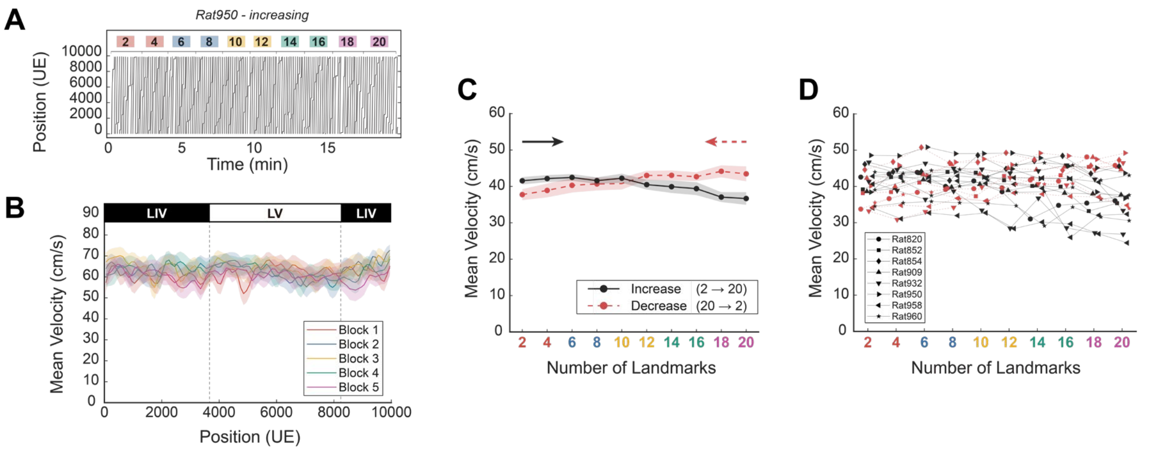


**Figure S1. Behavioral performance across parametric landmark accumulation conditions (Related to Figure 1).**

(A) Representative behavioral session. Time-position plot from a single representative session (Rat 950, landmark-increasing condition). The colored bars above the plot indicate the 10 sequential landmark conditions (from 2 to 20 landmarks). The rat maintained consistent running behavior throughout the VR navigation.

(B) Mean velocity profile along the virtual track. Population mean velocity (n = 8 rats, all sessions) plotted against track position across the five blocks. Velocity remained stable regardless of the position within the track (LV vs LIV zones) or the cumulative number of landmarks. Linear Mixed-Effects (LME) analysis, treating individual rats as random effects, confirmed no significant difference in velocity across blocks ($F(1, 108) = 0.45$, $p = 0.51$), indicating that the parametric manipulation did not induce systematic changes in running speed.

(C) Independence of running velocity from landmark density. Mean velocity plotted as a function of the sequential condition order within a session for both increasing (black solid line, 2-to-20 landmarks) and decreasing (red dashed line, 20-to-2 landmarks) sessions. The overlapping profiles demonstrate that running speed is independent of the absolute number of landmarks presented on the track. LME analysis confirmed that while a minor decrease in velocity occurred due to time-dependent factors ($F(1, 225) = 44.97$, $p < 0.001$), the number of landmarks had no significant impact on locomotor behavior ($F(1, 225) = 0.31$, $p = 0.58$). These results ensure that the observed subregional neural differences are not confounded by behavioral variations.

(D) Individual behavioral consistency. Mean velocity for individual rats and sessions across the 10 landmark conditions. Although baseline running speeds varied between individuals (accounted for as random intercepts in the LME model), all subjects exhibited a consistent behavioral trend across conditions. These results demonstrate that behavioral performance was robust and comparable across the entire animal cohort, regardless of the number of landmarks.

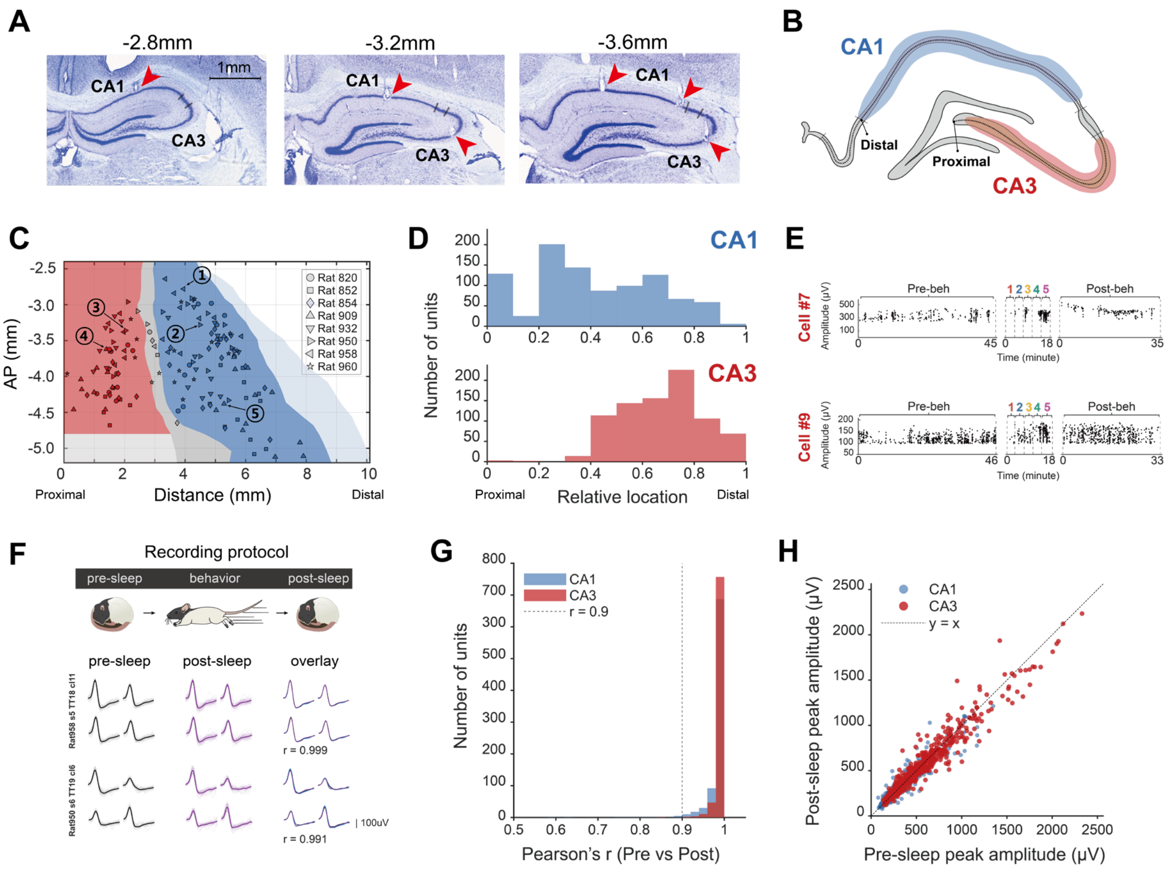


**Figure S2. Histological verification of recording sites and recording stability (Related to Figure 1).**

(A) Histological verification of tetrode locations. Nissl-stained sections show electrode tracks terminating in the CA1 and CA3 pyramidal layers. Red arrowheads indicate representative tetrode tips. (Scale bar, 1 mm).

(B) Schematic of the hippocampal proximodistal axis. Diagram illustrating the anatomical boundaries of CA1 (blue) and CA3 (red) subregions. The longitudinal axis is linearized to define the relative position from the proximal (dentate gyrus junction) to the distal end.

(C) Linearized flat map of recording sites across the proximodistal axis. Plot showing the tetrode tip locations from all rats (n = 8) projected onto a standardized flat map coordinate system (AP coordinate vs. proximodistal distance). Different markers represent individual animals, confirming broad and consistent coverage of both CA1 and CA3 subregions.

(D) Normalized distribution of recorded units. Histograms showing the number of units recorded along the normalized proximodistal axis (0: Proximal, 1: Distal). While recording sites covered the proximodistal extent of CA1 (top), CA3 recordings (bottom) were intentionally focused toward the intermediate and distal segments to capture representative spatial dynamics of the autoassociative network.

(E) Recording stability assessment. Spike amplitudes of representative CA3 units (Cell #7 and #9) are plotted over time, including pre-behavioral sleep, the behavioral epoch, and post-behavioral sleep. Stable spike amplitudes during sleep sessions confirm that the quiescence observed during behavioral blocks in CA3 reflects functional silence rather than recording instability or loss of the unit.

(F) Recording protocol and representative waveform stability. (Top) Schematic of the experimental timeline consisting of pre-behavioral sleep, behavioral navigation in VR, and post-behavioral sleep. (Bottom) Average spike waveforms from representative CA1 and CA3 units recorded during pre-behavioral sleep (black) and post-behavioral sleep (purple). High waveform similarity between epochs indicates minimal electrode drift.

(G) Population-level waveform correlation. Distribution of Pearson's correlation coefficients (r) between pre- and post-behavior sleep average waveforms for all valid units (CA1: 904, CA3: 815). The vast majority of units (98.8%) maintained high stability with $r > 0.90$ (Mean $r = 0.99$, Median $r > 0.99$), confirming that the identity of single units was preserved throughout the duration of the experiment.

(H) Spike amplitude stability. Comparison of peak-to-valley spike amplitudes between pre- and post-behavior sleep sessions. No significant difference was observed in mean amplitudes across sessions ($Z = 0.83$, $p = 0.40$; Wilcoxon signed-rank test). This robust stability ensures that changes in firing rates during behavior (e.g., CA3 quiescence in sparse environments) are functional signatures rather than technical artifacts of recording stability.

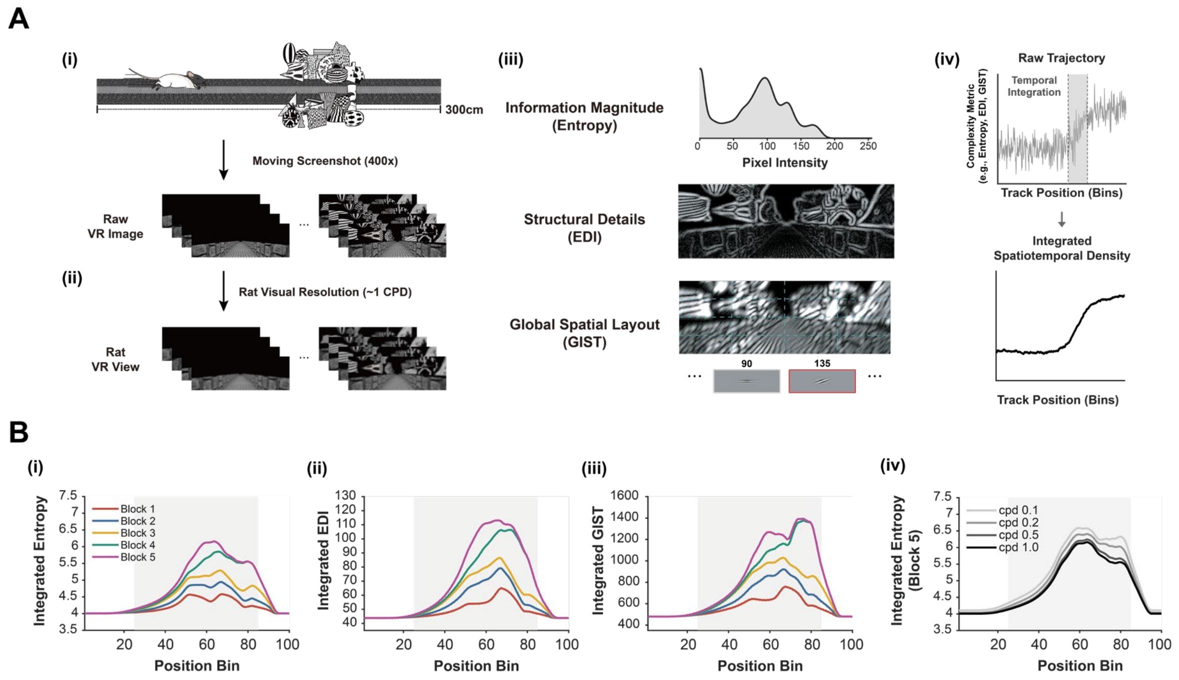


**Figure S3. Quantification of multidimensional visual scene complexity and spatiotemporal integration (Related to Figure 3).**

(A) Computational pipeline for extracting continuous visual features from the virtual (VR) environment. (i) Schematic top-down view of the VR track, illustrating the most complex landmark accumulation condition (Block 5). To continuously quantify the visual input during navigation, the 300-cm virtual track was uniformly divided into 400 spatial bins, and raw VR screenshots were extracted at each discrete position. (ii) To closely model the biological constraints of the rodent visual system, raw images were downsampled to match the visual acuity of Long-Evans rats, estimated at approximately 1.0 cycle per degree (CPD)[1]. This step ensures that the subsequent complexity analysis reflects the actual visual information perceived by the animals, utilizing the biological visual resolution of the rat rather than the high resolution of the monitors. (iii) The biologically constrained images (resized to 512 × 512 pixels) were analyzed using three distinct metrics to independently quantify different dimensions of visual complexity. Information Magnitude (Entropy): Shannon's entropy[2] was calculated to measure the statistical dispersion of grayscale pixel intensities, representing the overall richness and local texture of the scene. Structural Details (EDI): The Edge Density Index was computed by extracting gradient magnitudes using a Sobel filter[3,4], quantifying the density and strength of sharp artificial boundaries and local contours. Global Spatial Layout (GIST): The macroscopic geometric layout was evaluated using a Gabor filter bank[5] consisting of three spatial scales (wavelengths: 4, 8, and 16) and four orientations (0°, 45°, 90°, and 135°), capturing the spatial envelope and large-scale structural energy of the environment. (iv) To model the biological accumulation of visual evidence necessary for hippocampal network recruitment, the instantaneous feature values were spatiotemporally integrated. A causal trailing moving average was applied, where the spatiotemporal density at a given bin n was computed by averaging the visual inputs strictly from the past 45 bins (i.e., from bin n-44 to bin n). This approach translates discrete spatial snapshots into a continuous, accumulated sensory drive.

(B) Spatiotemporal dynamics of visual scene complexity across the parametric landmark accumulation paradigm. In all subpanels, the gray shaded area indicates the Landmark Visible (LV) zone. Integrated spatiotemporal trajectories of (i) Information Magnitude (Entropy), (ii) Structural Details (EDI), and (iii) Global Spatial Layout (GIST), respectively, across the 100 position bins. Color-coded lines correspond to the five blocks. Across all three metrics, the integrated visual complexity exhibits a continuous and gradual buildup as the animal navigates through the LV zone, parametrically scaling from Block 1 to Block 5. This continuous, linear accumulation of visual sensory drive highlights a critical dissociation: while the environmental input increases progressively, the hippocampal CA3 network exhibits an abrupt, nonlinear increase in place cell recruitment, supporting a threshold-gated map formation mechanism. (iv) Robustness of the complexity trajectory across varying biological visual acuities. The integrated entropy for the most complex condition (Block 5) was recalculated using four different spatial resolutions (CPD: 0.1, 0.2, 0.5, and 1.0). Although absolute values shift slightly depending on the resolution, the overarching spatiotemporal trend remains consistent, confirming that the observed continuous sensory buildup is a robust environmental property rather than an artifact of a specific downsampling parameter.

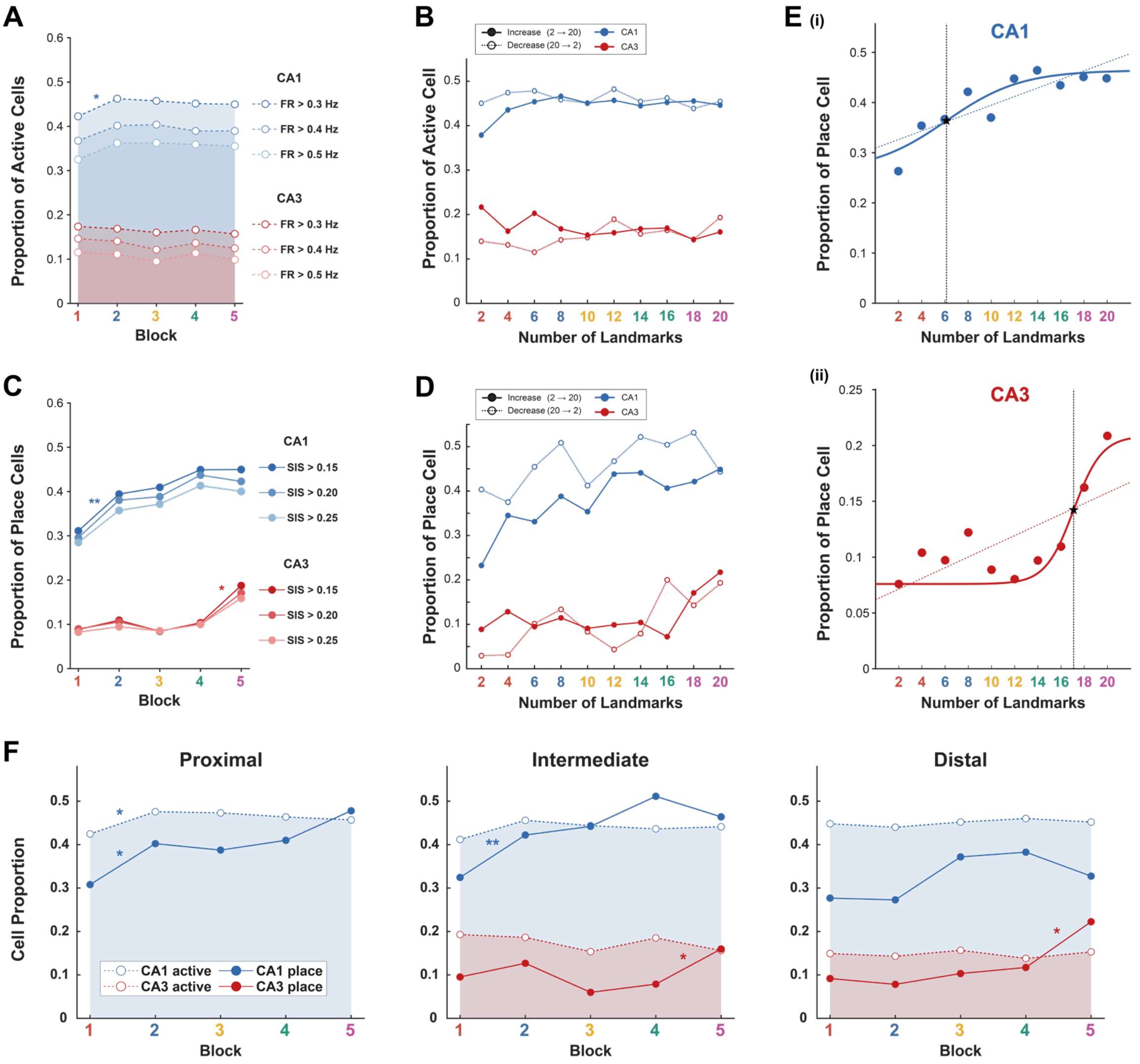


**Figure S4. Robustness of hippocampal spatial map recruitment across analytical criteria, manipulation sequence, and the proximodistal axis (Related to Figure 3).**

(A) Sensitivity of active cell recruitment to firing rate (FR) thresholds. Proportions of active cells across the five blocks, defined by various FR thresholds (FR > 0.3, 0.4 and 0.5 Hz). In both CA1 and CA3, active cell proportions remained stable across blocks regardless of the stringency of the firing rate criterion, indicating that the overall recruitment levels are not biased by active cell threshold.

(B) Proportion of active cells across experimental conditions. The fraction of active cells remained stable across all 10 landmark conditions in both CA1 (blue) and CA3 (red), regardless of the sequence direction (Increase: solid lines; Decrease: dotted lines). This stability indicates that global network excitability was not biased by the order of landmark presentation during the parametric manipulation.

(C) Sensitivity of place cell recruitment to spatial information (SI) thresholds. Proportions of units identified as place cells using different SI criteria (SI > 0.15, 0.20, and 0.25 bits/spike). In CA1, place cell proportions showed early recruitment and a gradual increase across blocks. In CA3, the characteristic nonlinear increase in spatial representation at Block 5 was robustly observed across all tested SI thresholds (Block 4 vs. 5 in CA3: all $p < 0.05$, $\chi^2$ test), demonstrating that the spatial map recruitment is independent of spatial selectivity criteria.

(D) Consistency of place cell recruitment dynamics. Proportion of units identified as place cells across the 10 conditions. To validate the consistency of spatial map formation across different landmark manipulation sequences, recruitment trajectories were analyzed using a nonlinear logistic regression model. This model incorporated both linear and quadratic terms ($Conditions^2$) to capture the characteristic sigmoidal surge in CA3 spatial representation. In both subregions, the recruitment profiles were statistically indistinguishable between the increasing and decreasing sequences, as evidenced by the lack of significant interactions between condition terms and sequence direction (all interaction p-values > 0.25). Notably, the nonlinear increase in CA3 recruitment was robustly observed in both sequences, supporting the pooling of these datasets for population-level analyses. While the elevated proportion of CA3 place cells persisted for one additional condition during the decreasing sequences (i.e., 16 landmarks; Increase vs. Decrease: $\chi^2(1) = 4.75$, $p = 0.029$), transition-focused GLMs confirmed that the rate of state change (formation vs. collapse) did not significantly differ between sequences ($p > 0.08$ for all adjacent transitions). This subtle divergence likely reflects the inherent attractor properties of the CA3 network (i.e., hysteresis), further confirming that the observed recruitment is a stable, environment-driven regional property.

(E) Population recruitment data for (i) CA1 and (ii) CA3 across the 10 landmark conditions were fitted to linear (dotted lines) and bounded sigmoidal models (solid lines). Following established methodologies[6], the sigmoidal model was constrained by empirical minimum and maximum asymptotes to standardize the degrees of freedom for a rigorous Akaike Information Criterion (AIC) comparison. While map recruitment in CA1 was better explained by the sigmoidal model over the linear fit ($\Delta$ AIC = 4.53), it exhibited an early inflection (~6 landmarks) with a relatively gradual slope (steepness, $k = 0.31$). In contrast, the recruitment trajectory in CA3 was strongly explained by the non-linear sigmoidal model ($\Delta$ AIC = 7.30), demonstrating a late inflection (~17 landmarks) and a steep transition ($k = 0.82$) compared to CA1. The inflection points of the sigmoidal fits are indicated by vertical dashed lines. This quantitative model superiority validates that the CA3 network undergoes a delayed, nonlinear state transition rather than a continuous integration of sensory input.

(F) Proportions of active cells (dotted lines) and place cells (solid lines) across five blocks in the proximal (left), intermediate (middle), and distal (right) zones of CA1 (blue) and CA3 (red). The proximodistal axis was defined by dividing the normalized coordinates into three equal segments (1/3 each). Note that the proximal CA3 subregion was excluded from this analysis due to an insufficient sample size. Overall, all analyzed subregions along the proximodistal axis exhibited a similar trend of spatial map recruitment in response to the increasing number of visual landmarks. Pairwise comparisons revealed significant increases in cell proportions at specific block transitions (CA1 proximal: active, Block 1 vs. 2: $\chi^2(1) = 3.93$, $p = 0.047$; place, Block 1 vs. 2: $\chi^2(1) = 6.45$, $p = 0.011$; CA1 intermediate place, Block 1 vs. 2: $\chi^2(1) = 7.18$, $p = 0.007$; CA3 intermediate place, Block 4 vs. 5: $\chi^2(1) = 4.13$, $p = 0.042$; CA3 distal place, Block 4 vs. 5: $\chi^2(1) = 4.20$, $p = 0.041$; $\chi^2$ tests).

* $p < 0.05$, ** $p < 0.01$

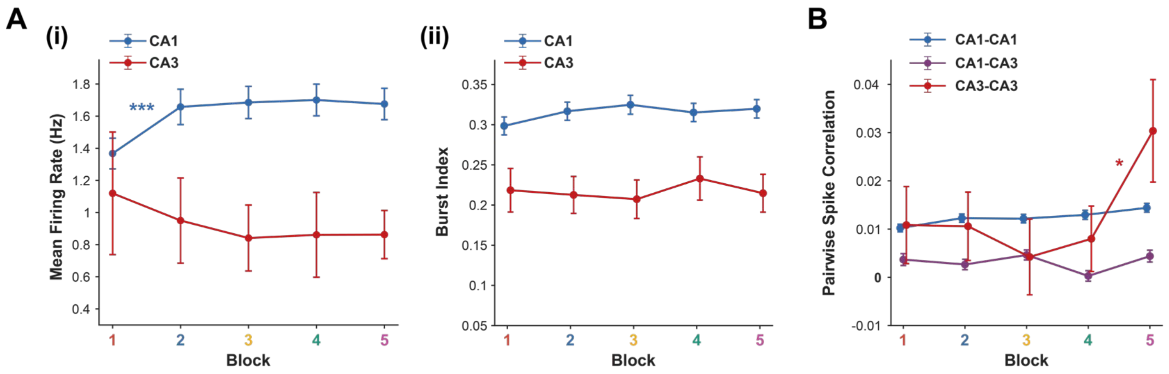


**Figure S5. Single-unit firing properties and network synchronization during spatial map recruitment (Related to Figure 3).**

(A) Firing properties of place cells longitudinally tracked from Block 5. (i) Mean firing rate across blocks. CA3 (red) remained stable ($\chi^2(4) = 2.32$, $p = 0.68$; Kruskal-Wallis test), whereas CA1 (blue) increased modestly after Block 1 ($\chi^2(4) = 19.68$, $p = 0.0006$). The same stability was observed for the active population of each block (CA1: $\chi^2(4) = 5.11$, $p = 0.28$; CA3: $\chi^2(4) = 4.77$, $p = 0.31$). (ii) Burst index (proportion of spikes with inter-spike intervals < 6 ms). Both CA1 and CA3 maintained stable single-unit bursting properties across all blocks (CA1: $\chi^2(4) = 2.47$, $p = 0.65$; CA3: $\chi^2(4) = 0.39$, $p = 0.98$). Together, these indicate that neither the amount nor the mode of single-unit firing changed with spatial map recruitment. Data are presented as means ± SEM.

(B) Mean pairwise spike correlations (100-ms bins) among place cells identified in Block 5 and tracked backward through the preceding blocks, as in Figure 3C. While CA1 intra-regional (CA1-CA1) and inter-regional (CA1-CA3) synchrony remained relatively constant, CA3 intra-regional synchrony (CA3-CA3) exhibited an abrupt increase at Block 5 (Block 1-4 vs. Block 5, $p = 0.013$; 10,000-iteration permutation test). This demonstrates that the recruitment of the CA3 spatial map is driven by synchronous network engagement. Data are presented as means ± SEM.

* $p < 0.05$, *** $p < 0.001$

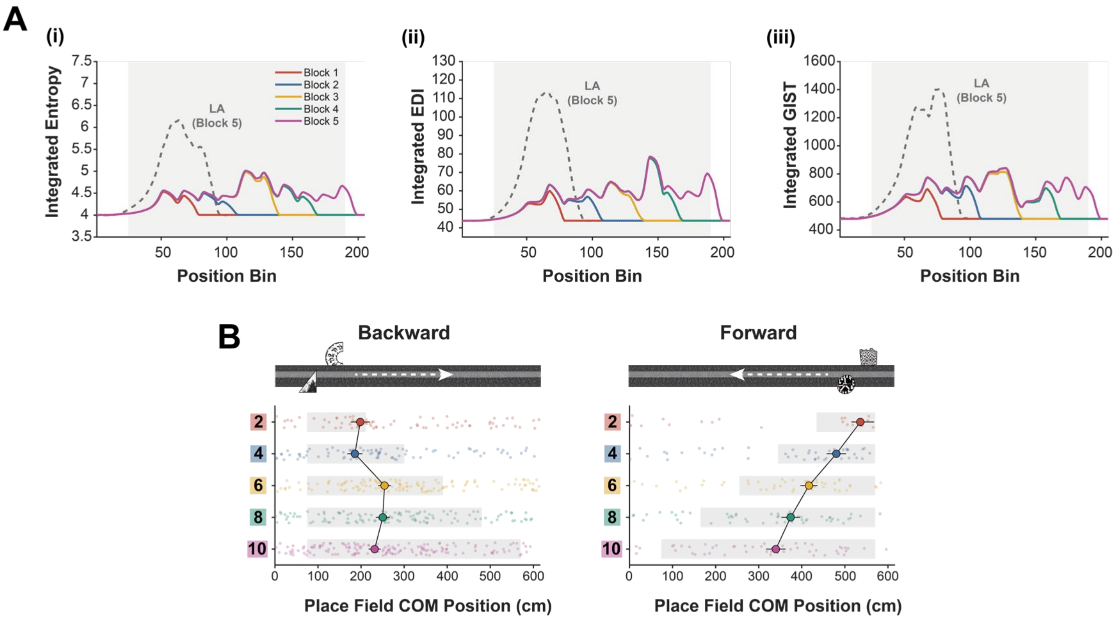


**Figure S6. Population dynamics and continuous integration of the CA1 spatial map in the Distributed Landmark paradigm (Related to Figure 4).**

(A) Integrated spatiotemporal trajectories of (i) Entropy, (ii) EDI, and (iii) GIST across the position bins. Color-coded solid lines correspond to the five blocks in the Distributed Landmarks (DL) paradigm. Y-axes are scaled identically to the Landmark Accumulation (LA) paradigm (Figure S3B). The dashed gray line indicates the reference complexity trajectory from Block 5 of the LA paradigm. The gray shaded area represents the LV zone of Block 5 in the DL paradigm. Unlike the continuous buildup in the LA paradigm, distributing landmarks prevents the formation of a dense ensemble. Consequently, visual complexity across all metrics remains persistently low throughout the environment.

(B) Continuous spatial updating of the CA1 population map in response to incremental landmark presentation. CA3 is excluded from this analysis because very few place cells were recruited in the DL paradigm. In this paradigm, landmarks were incrementally presented without forming a dense visual ensemble, with the presentation progressing either in the forward or backward direction (top schematics). (Bottom) Circular center of mass (COM) of individual CA1 place fields (dots) and the population circular mean ± SEM across the 600-cm virtual track. Gray shaded regions indicate the landmark-visible (LV) zone for each block. To accurately quantify continuous population map shifts across the teleportation boundary (0/600 cm), unwrapped circular COMs were analyzed using a linear regression model. The CA1 population COM shifted significantly opposite to the direction of landmark presentation — forward during backward-direction sessions (slope = 11.66 cm/block, $R^2 = 0.01$, $p = 0.018$; linear regression) and backward during forward-direction sessions (slope = -54.38 cm/block, $R^2 = 0.19$, $p < 0.001$). These results demonstrate that CA1 continuously integrates newly presented landmarks, dynamically updating its spatial map in accordance with environmental changes.

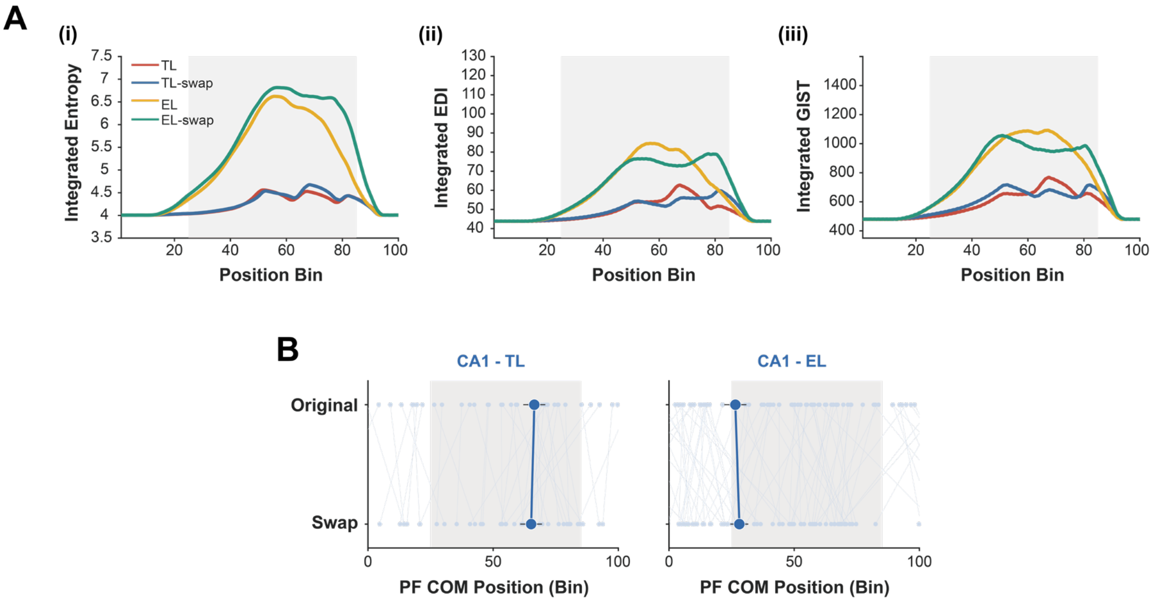


**Figure S7. Single-unit representational dynamics in the Landmark Swap paradigm (Related to Figure 5).**

(A) Integrated spatiotemporal trajectories of (i) Entropy, (ii) EDI, and (iii) GIST across the position bins. Color-coded solid lines correspond to the four conditions in the Landmark Swap paradigm: Triple Landmarks (TL), TL-swap, Ensemble Landmarks (EL), and EL-swap. The gray shaded area represents the LV zone. Consistent with the Landmark Accumulation paradigm, the presentation of a landmark ensemble (EL and EL-swap) generated substantially higher visual complexity across all metrics compared to the sparse landmarks (TL and TL-swap).

(B) Center of mass (COM) shifts of place fields in CA1 across original and swap conditions. Thin lines and dots represent individual cells, while thick solid lines indicate the circular mean ± SEM COM shift. The gray shaded region denotes the LV zone. To evaluate spatial displacement, only cells classified as place cells in both the original and swap configurations were included (CA1: n = 39 for TL, 69 for EL). Single-unit COM tracking in CA1 reveals local spatial shifts between the two configurations. CA3 was excluded from this visualization due to a near-complete absence of cells maintaining place fields across both configurations (n = 1 for TL, n = 0 for EL), consistent with its global remapping.

*** $p < 0.001$

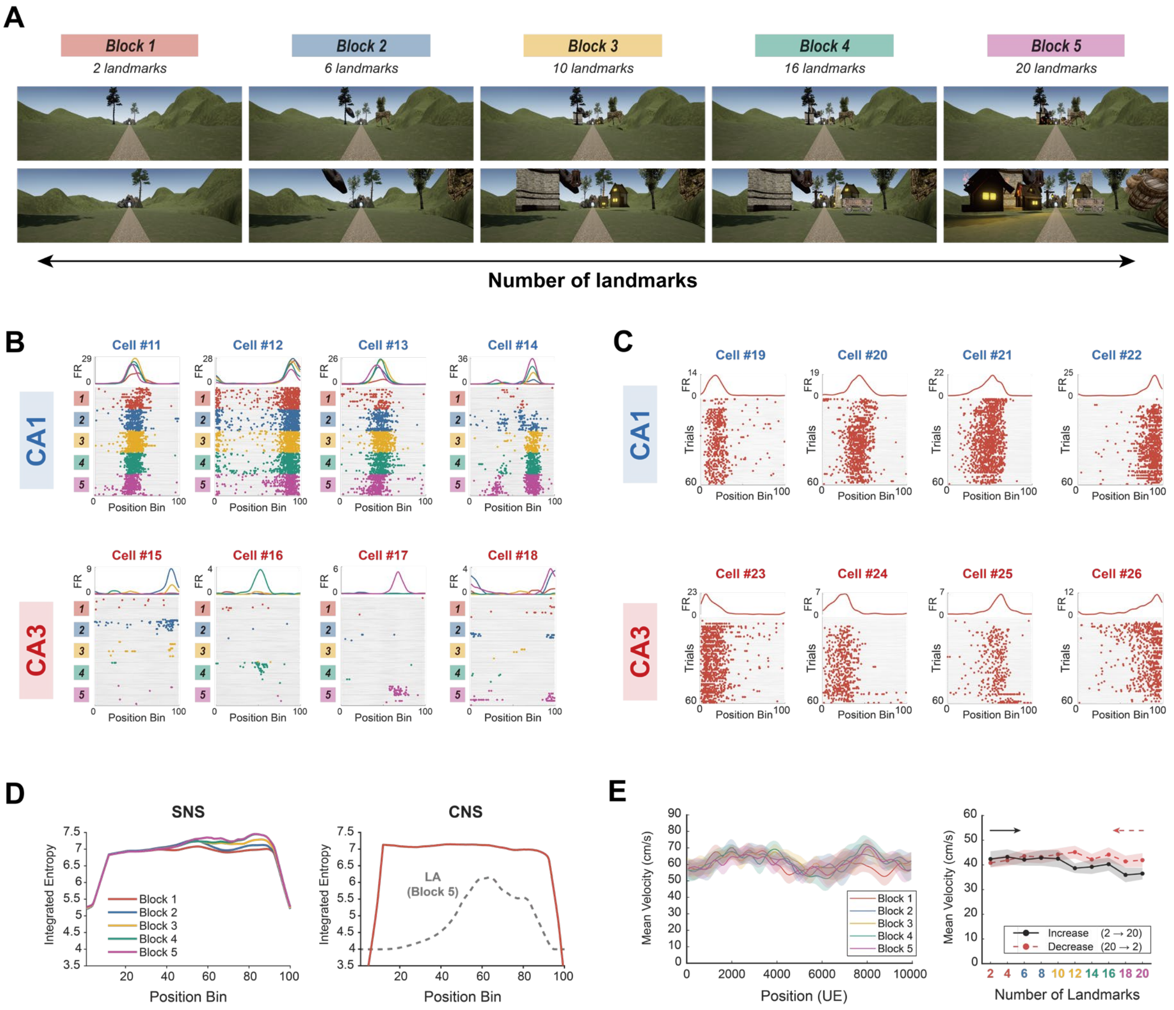


**Figure S8. The naturalistic environments elicit robust place coding while visual complexity and running behavior remain controlled (Related to Figure 6).**

(A) Full sequence of the Simple Natural Scene (SNS) accumulation protocol. Representative views of the SNS environment across all five blocks (Blocks 1–5), in which naturalistic landmarks were incrementally scaled from 2 to 20 within each session in both ascending and descending sequences (from 2 to 20 landmarks) against a continuous natural background, following the same parametric accumulation protocol used in the visually Poor environment (Figure 1C). The background scene was held constant while landmark density increased, confirming that the manipulation reflects landmark accumulation rather than a change of environment.

(B) Representative responses of CA1 (Cell #11-14) and CA3 (Cell #15-18) place cells across 5 blocks in the SNS environment.

(C) Representative responses of CA1 (Cell #19-22) and CA3 (Cell #23-26) place cells in the CNS environment.

(D) Quantification of visual scene entropy. Integrated entropy across the track for (Left) the five sequential blocks (colored lines) in the Simple Natural Scene (SNS) paradigm and (Right) the single Complex Natural Scene (CNS) condition (red line). For comparison, the dashed gray line indicates Block 5 of the Landmark Accumulation paradigm in the Poor environment. The sharp drop in entropy at both ends of the track is due to a tunnel structure designed for natural teleportation.

(E) Behavioral performance in the SNS paradigm. (Left) Population mean velocity plotted against track position for Blocks 1-5. Running speed remained stable with no significant difference across blocks ($F(1, 58) = 0.22$, $p = 0.64$; Linear Mixed-Effects model). (Right) Mean velocity across the 10 sequential conditions (increasing and decreasing). Within-session running speed declined modestly with time-on-task rather than with the number of landmarks, which varied oppositely with time in the increasing and decreasing sequences; this behavioral stability is comparable to the Landmark Accumulation paradigm (Figure S1). Behavioral performance in the CNS environment was reported in a previous study[7].

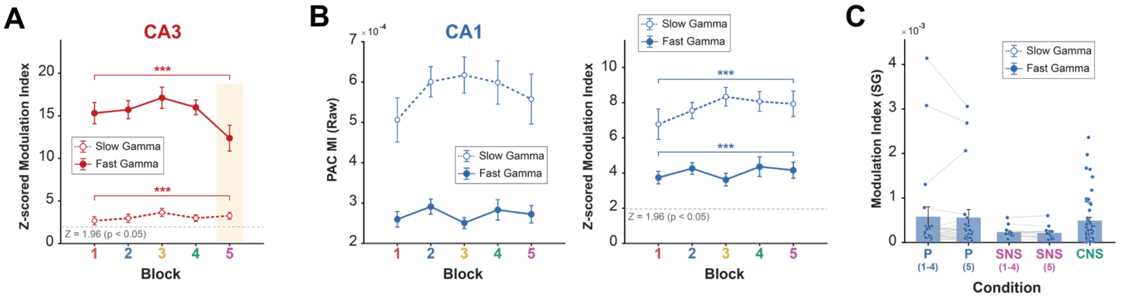


**Figure S9. Statistical validation and subregional dissociation of PAC dynamics (Related to Figure 7).**

(A) Surrogate Z-scored modulation index (MI) for fast gamma (FG, solid line) and slow gamma (SG, dashed line) in CA3 across the five blocks of the Landmark Accumulation paradigm. Z-scores were calculated against surrogate null distributions generated by randomly shifting the amplitude time series relative to the theta phase ($n = 200$ iterations). The dashed gray line denotes $Z = 1.96$, which corresponds to the threshold for single-session statistical significance ($p < 0.05$). The orange shaded area indicates Block 5, where the nonlinear state transition of the CA3 spatial map occurs. Both FG and SG PAC levels were significantly greater than chance level ($Z = 0$) across all blocks (all $p < 0.001$; one-sample t-test). Furthermore, the nonlinear reduction in FG-PAC at Block 5 was statistically significant within the Z-scored domain (Block 1-4 vs. Block 5: $\beta = -3.66$, $p = 0.0067$; linear mixed-effects model), confirming that the observed decoupling represents a physiological state change rather than a signal artifact. Data are presented as means ± within-subject SEM.

(B) Distinct PAC dynamics in CA1. In contrast to the nonlinear sensory decoupling observed in CA3, CA1 exhibited stable PAC levels across all landmark conditions. The left panel displays the raw MI, and the right panel displays the surrogate Z-scored MI for both FG (solid blue line) and SG (dashed blue line). Both metrics remained constant throughout the parametric accumulation of visual landmarks, as confirmed by statistical comparisons between Block 1-4 and Block 5 (Raw MI: all $p > 0.68$; Z-scored MI: all $p > 0.72$; linear mixed-effects model) and between early stages (Block 1 vs. Block 2: $Z = 0.67$, $p = 0.51$; Wilcoxon signed-rank test). Error bars indicate within-subject SEM.

(C) Modulation index of SG in CA1 across the five stages of scene complexity. In contrast to the substantial elevation of SG coupling observed in CA3 within the CNS environment (Figure 7G), CA1 SG-PAC remained relatively stable and did not exhibit distinct network-level transitions across the different environmental complexities (Poor Block 5 vs. CNS: $Z = -0.34$, $p = 0.74$; Mann-Whitney U test). Bars and error bars represent the mean and the between-subject SEM. Single dots indicate individual sessions, and gray lines connect paired sessions.

*** $p < 0.001$

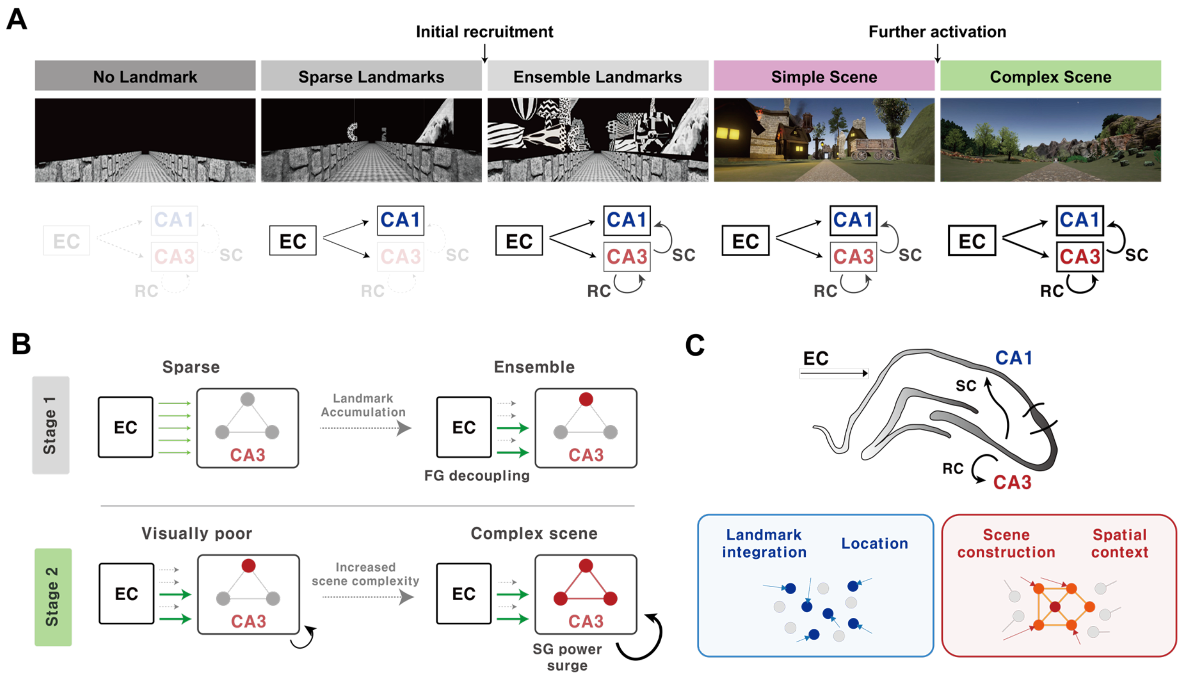


**Figure S10. Conceptual model for hippocampal map recruitment: CA3 scene construction vs. CA1 continuous integration**

(A) Schematic of network state transitions across a gradient of visual complexity. In visually "Poor" environments, the CA3 autoassociative network remains quiescent, while CA1 initializes a provisional spatial map. Increasing scene complexity—via visual ensembles or background scaffolds—triggers a threshold-gated state transition in CA3, leading to the de novo recruitment of a coherent spatial map. RC, recurrent collaterals; SC, Schaffer collaterals; EC, entorhinal cortex.

(B) Stepwise trajectory of CA3 map recruitment. Stage 1: Upon reaching the visual ensemble threshold, localized subnetwork activation triggers potent disynaptic feedback inhibition, effectively decoupling the network from subsequent feedforward sensory drives (FG-PAC attenuation). Stage 2: In a highly dense visual context, convergent excitatory drive reaches the critical mass of the recurrent network. This shifts the excitation-inhibition balance toward sustained recurrent excitation, triggering global network engagement and recurrent synchronization (SG-PAC surge).

(C) Conceptual framework for the subregional division of labor. Scene construction in CA3 (Red): CA3 acts as a nonlinearly gated 'scene constructor' that validates relational coherence to anchor the cognitive map to a stable, holistic spatial context (attractor state). Landmark integration in CA1 (Blue): CA1 serves as a 'continuous integrator' that linearly accumulates landmarks and compares incoming sensory evidence (EC) with scene-based predictions from CA3. Together, this synergy balances representational specificity with navigational flexibility.
